# Electrochemical cofactor recycling of bacterial microcompartments

**DOI:** 10.1101/2024.07.15.603600

**Authors:** Markus Sutter, Lisa M. Utschig, Jens Niklas, Sathi Paul, Darren N. Kahan, Sayan Gupta, Oleg G. Poluektov, Bryan H. Ferlez, Nicholas M. Tefft, Michaela A. TerAvest, David P. Hickey, Josh V. Vermaas, Corie Y. Ralston, Cheryl A. Kerfeld

**Author notes:** Department of Molecular Biology and Genetics, Cornell University; Ithaca, NY 14583, USA.

## Abstract

Bacterial microcompartments (BMCs) are prokaryotic organelles that consist of a protein shell which sequesters metabolic reactions in its interior. While most of the substrates and products are relatively small and can permeate the shell, many of the encapsulated enzymes require cofactors that must be regenerated inside. We have analyzed the occurrence of an enzyme previously assigned as a cobalamin (vitamin B_12_) reductase and, curiously, found it in many unrelated BMC types that do not employ B_12_ cofactors. We propose NAD+ regeneration as a new function of this enzyme and name it MNdh, for Metabolosome NADH dehydrogenase. Its partner shell protein BMC-T^SE^ assists in passing the generated electrons to the outside. We support this hypothesis with bioinformatic analysis, functional assays, EPR spectroscopy, protein voltammetry and structural modeling verified with X-ray footprinting. This discovery represents a new paradigm for the BMC field, identifying a new, widely occurring route for cofactor recycling and a new function for the shell as separating redox environments.

## Introduction

Bacterial microcompartments are protein shells encapsulating enzymes that are found in a large variety of prokaryotes^1,2^. There are two distinct major classes, anabolic carboxysomes found mostly in cyanobacteria that fix carbon dioxide with encapsulated RuBisCO^3^, and catabolic metabolosomes that process a variety of organic compounds^4^. The canonical reaction mechanism of metabolosomes starts with an encapsulated signature enzyme that generates an aldehyde that is then processed by an aldehyde dehydrogenase to a Coenzyme A adduct (Fig. 1A)^2,4,5^. The aldehyde dehydrogenase reaction involves a reduction of NAD+ to NADH, and the regeneration of NAD+ is thought to proceed through a side reaction that generates an alcohol from the aldehyde substrate (Fig. 1A)^6^. The most well-known metabolosomes are the EUT (ethanolamine degradation)^7^ and PDU (propanediol degradation)^8^ BMCs. More recently a number of additional metabolosomes have been characterized: PVM (Planctomycete and Verrucomicrobia microcompartment^9^, GRM (Glycyl radical BMCs)^10,11^, aminoethane degrading (RMM: Rhodococcus and Mycobacteria microcompartment or AAUM: aminoethane utilizing microcompartment)^12^, a taurine degrading^13,14^ and an aromatic substrate degrading BMC (ARO)^15^.

**Figure 1.**
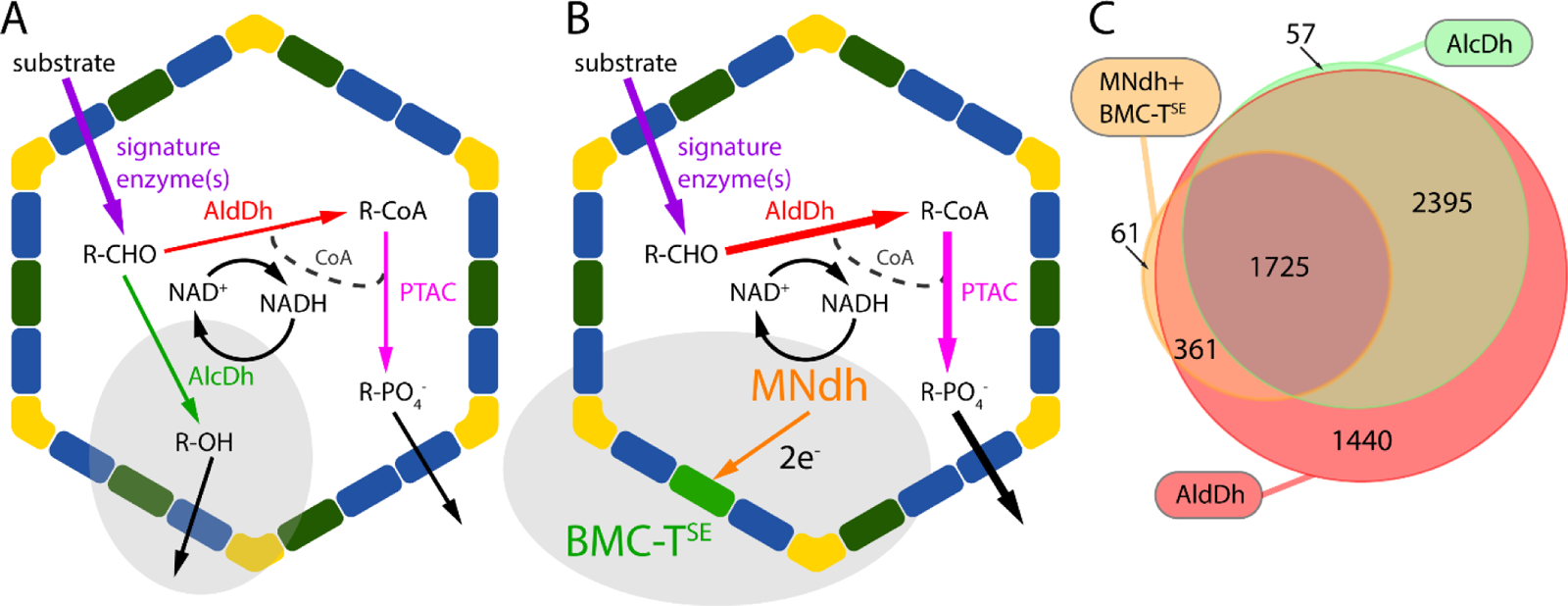
Metabolosome reaction schemes and MNdh distribution. (**A**) Traditional metabolosome scheme. (**B**) MNdh NAD+ regeneration scheme. (**C**) Co-occurrence of NAD+ cofactor regeneration enzymes with AldDh in BMC loci shown as an Euler diagram (not visualized is an overlap of AlcDh and MNdh/BMC-TSE and no AldDh with 25 occurrences). Abbreviations: AldDh: Aldehyde dehydrogenase; AlcDh: Alcohol dehydrogenase; PTAC: phosphotransacetylase; CoA: Coenzyme A.

Metabolosome BMCs are typically between 55-200 nm in diameter^8,9^ and their shells consist of proteins formed from two structural domains that are conserved across all BMCs. The BMC-H and BMC-T contain the pfam00936 domain and form hexameric and trimeric hexagonal tiles, respectively. The hexagonal tiles have a central pore at the center of symmetry that is thought to be the main conduit into the shell for metabolites^16^; however their typical diameter of around 5 Å make them less suitable for larger molecules like cofactors. For the most common type of BMC-T, the BMC-T^S^, which consist of a tandem fusion of two pfam00936 domains that do not further dimerize (hence superscript “S” for single), there are two main subtypes on a phylogenetic tree^1^: the carboxysome shell protein CcmO and proteins related to PduT, a BMC-T^S^ first characterized from the PDU BMC^17^ which contains a 4Fe-4S cluster in its central pore. However, there are many BMC-T proteins that do not have the pore motif (CPGK, sometimes CSGK) for the metal cluster binding at the pore. The second structural domain necessary for the formation of BMC shells is the pfam03319 domain that forms the BMC-P pentamers^18^ that cap an overall icosahedral or polyhedral structure^19–23^.

When looking at model shells that contain representative of shell protein components^19^ it seems unlikely and inefficient that larger cofactors such as NADH or Coenzyme A are exchanged across the shell for each turnover. Moreover, pores in the shell protein tiles are typically too small to permit passage of cofactor molecules. There are examples of gated pores of larger diameter shell proteins (BMC-T^D^)^19,24–26^ that could conceivably conduct cofactors, however these are not found in all shells, they typically are a minor fraction when present^27^. This would be relatively low throughput and would require a concentration gradient to be established across the shell. Instead, it is more likely that cofactors are acquired during assembly (as in the case of the CoA associated with the PTAC, PduL^28^) and regenerated by core metabolosome enzymes. As such they form a private cofactor pool for organelle function.

As part of a previous bioinformatics analysis^1^ we surveyed the distribution of all enzymes within predicted BMC loci and noticed that the enzyme known as PduS is found frequently in loci other than the PDU; indeed it is distributed across many functionally diverse BMC types and species. The proposed function of this protein in literature is performing the reduction from aquacob(III)alamin to cob(I)alamin as part of the regeneration of inactivated adenosylcobalamin^29–31^ in Vitamin B_12_/cobalamin-dependent metabolosomes. This specialized function, however, is unlikely to be relevant in many of the non-PDU or EUT BMC types as there are no known BMC related enzymes using B_12_ other than PDU and EUT signature enzymes. Furthermore, we found that consistently adjacent to it there was a PduT homolog that contained the motif for FeS cluster binding. Here we present a comprehensive bioinformatic analysis of this noncanonically occurring PduS/PduT pair, structural modelling and functional and spectroscopic data to show that the PduS-like protein acts in conjunction with its cognate PduT-like protein to pass two electrons from the FMN cofactor via three 4Fe-4S clusters to an acceptor outside the shell, thereby regenerating NAD+ within the BMC. Accordingly, we propose a new, more universal name for PduS, MNdh, for Metabolosome NADH dehydrogenase, and we rename its co-occurring PduT homolog BMC-T^SE^ (E for electron transfer) because it contains a central 4Fe-4S cluster. BMC-T^SE^ and MNdh form a cofactor regeneration system widespread across functionally diverse BMCs that highlights its importance. Knowledge of this housekeeping cofactor recycling module allows us to better understand the native metabolosome reaction mechanisms as well as enable the use of NAD recycling for engineered systems.

## Results

### Bioinformatics analysis of the MNdh / BMC-T^SE^ prevalence

We undertook a comprehensive survey for the presence of MNdh and for BMC-T^SE^ in all BMC loci, using the BMC locus dataset from Sutter *et al*^1^. The genes for those two proteins are almost exclusively located right next to each other and are observed in 2172 of over 7000 BMC loci, distributed across a large functional variety of metabolosomes (Table S1, Fig. S1). However, in conflict with the originally proposed function of the MNdh for Vitamin B_12_/cobalamin regeneration, many of the functionally varied BMC types are not predicted to utilize B_12_. Among them are the more recently characterized GRM, PVM and Taurine utilizing BMC (also known as GRMguf)^9,11,13,14^ as well as many BMC types with functions that have yet to be established (MIC, MUF, FRAG, SPU)^1,4^ (Fig. S1, Table S1). MNdh is only found in BMC types that also contain an aldehyde dehydrogenase that is essential for the BMC function for the established metabolosome types and likely also for the yet-to-be-characterized ones. This type of aldehyde dehydrogenase uses NAD+ as an electron acceptor, generating NADH. Typically, the NAD recycling is carried out by the co-encapsulated AlcDH (Fig. 1A), but we noted a large number (361) of BMCs lacking a gene for AlcDH, but containing PduS/T (Fig. 1C). This led us to formulate a new hypothesis that the primary function of MNdh is to regenerate the NAD+ cofactor inside the BMC lumen (Fig. 1B), passing the electrons to the outside of the BMC via the Fe-S cluster of BMC-T^SE^. An advantage of this type of regeneration versus the traditional type of NAD+ regeneration with alcohol dehydrogenase is that it prevents the need to produce the alcohol side product. In this pathway all of the input substrate is converted to the more valuable CoA-product (Fig. 1B).

The occurrence of MNdh in BMC functional types is variable, however many of them have a higher than 90% occurrence for a given type (Table S1, Fig. S1). Many of them co-occur with alcohol dehydrogenase that performs the canonical NAD+ regeneration (Fig. 1C, Table S1), like in the case of the highly characterized PDU type BMC of *Salmonella enterica*. Both AlcDh and MNdh/BMC-T^SE^ almost exclusively co-occur with AldDh, highlighting their functional dependence. The large number of loci that contain both NAD+ regeneration systems is an indication of their functional importance in cofactor recycling. Potentially they can be either both active at the same time or active under different circumstances, such as electron acceptor availability.

Additional evidence of a more general, “housekeeping” role in metabolosome function (as opposed to a substrate-specific role) for MNdh-BMC-T^SE^ emerges from consideration of their phylogeny and BMC type distribution. We generated a phylogenetic tree of MNdh sequences and analyzed their correlation with BMC type and phylogenetic origin (Fig. S2). There is surprisingly little correlation between the BMC type and sequence of MNdh. For example, the MNdh found in the PDU and EUT BMC type BMCs are all over the MNdh tree (Fig. S2). This indicates that the function of MNdh is not tied to a specific BMC reaction pathway but more likely a very generic one. This is in stark contrast to a phylogenetic tree of the aldehyde dehydrogenase where a strong clustering of the sequences by BMC type is observed^1^. Even the correlation with phylogenetic origin is not very strong, only a few phyla have their MNdh sequences clustered in one part of the tree, e.g. for Planctomycetes, Verrucomicrobia and Elusimicrobia. This could indicate a frequent occurrence of horizontal gene transfer of the MNdh/BMC-TSE module and also supports our hypothesis of MNdh/BMC-T^SE^ having a function applicable to all AldDh containing BMC types. In some BMC types the MNdh/BMC-T^SE^ also occur more frequently in satellite loci^32^ than main loci (PVM: 18 out of 20 occurrences are in satellite and 5 out of 7 for ACI), which suggests that they are a module that can be used under certain environmental conditions, such as anaerobic growth or specific redox conditions. In this way MNdh/BMC-T^SE^ module contributes metabolic flexibility to a given BMC functional type.

### Structural modelling of MNdh and the MNdh-BMC-T^SE^ complex

We generated an AlphaFold2 model^33,34^ of MNdh based on the sequence of the *C. botulinum* homolog that shows a high level of confidence throughout (Fig. 2A). We were able to place the 4Fe-4S clusters manually into the AlphaFold2 apo structure and the FMN and NAD+ using homologous structures (Fig. 2B). There are four distinct structural domains of MNdh: 1) the “Complex 1-51kDa” pfam01512, 2) the SLBB (soluble ligand binding β-grasp) pfam10531, 3) the 4Fe-4S dicluster pfam13634, and 4) the RnfC sandwich hybrid domain pfam13375. As the name suggests, the pfam01512 domain is found in a protein of the respiratory complex 1 (e.g. NuoF in *E. coli*); it covers the main part of the FMN/NAD binding domain. The pfam10531 SLBB and pfam13375 RnfC are adjacent to the FMN/NAD binding pocket and the pfam13634 4Fe-4S dicluster domain is located above (Fig. 2C). The AlphaFold2 model of MNdh closely resembles a recent Cryo-EM structure of RnfC, a Na+-translocating ferredoxin: NAD+ reductase^35^ (Fig. 2D). One notable difference is a circular permutation of the pfam13375 domain that is found at the C-terminus of RnfC compared to the N-terminus of MNdh (Fig. 2CD). Separating the RnfC into two parts and aligning each separately (to overcome the limitation of alignment tools to only consider contiguous alignment) gives rmsd values of 0.9 Å over 54 C-α atoms for the pfam13375 domain and 1.2 Å over 228 C-α atoms for the rest of the protein (Fig. S3). The Fe-S clusters and the residues for the FMN/NADH binding site align very well (Fig. S3, inset). However, as expected, we see that a large number of differences between RnfC and MNdh are located at the interface region with their respective binding partners (Fig. 2CD, Fig. S3, inset).

**Figure 2.**
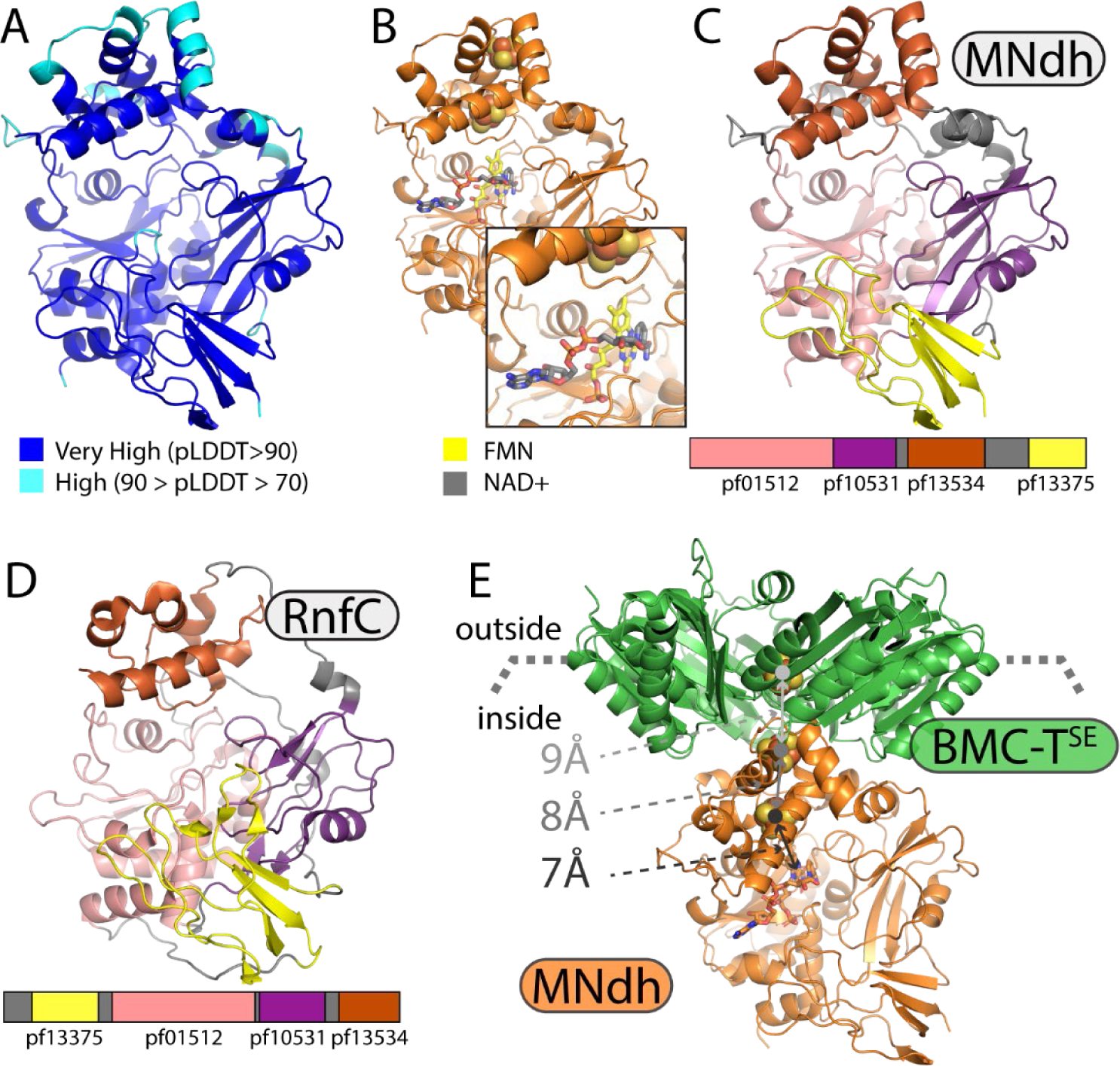
Structural models of MNdh. (**A**) AlphaFold2 confidence. (**B**) Cofactor and metal placement in the MNdh structure. (**C**) Domains of MNdh. (**D**) Domains of RnfC (**E**) Model of a MNdh-BMC-TSE complex.

We also generated an AlphaFold2^36^ prediction of a MNdh - BMC-T^SE^ complex (Fig. 2E). The solutions obtained are largely identical to each other, indicating a good convergence; however from 20 solutions, half of them predict binding to the exterior of the shell via BMC-T^SE^, the other half on the interior/lumenal surface (as in Fig. 2E). Physiologically, the convex-surface binding (MNdh inside the BMC) is the most plausible, not only based on the function of internal NAD recycling but also because MNdh has been found in isolated intact BMCs^13,37^; non-covalent association of MNdh to the external surface of a BMC is unlikely to persist through purification. Further circumstantial evidence for an internal localization comes from much higher amino acid conservation observed for residues on the inside, but only for the BMC-T^SE^ that co-occur with MNdh (Fig. S4). In the BMC-T^SE^-MNdh complex, the Fe-S cluster of BMC-T^SE^ is located along a straight line between the FMN/NADH binding site and the two Fe-S clusters of MNdh with only 7-9 Å between them (Fig. 2E), suggestive and of an electron transfer along this path analogous to the one seen in RnfC^35^.

### X-ray footprinting of the MNdh-BMC-T^SE^ complex

To experimentally validate the AlphaFold2 model of the complex, we performed X-ray footprinting mass spectrometry (XFMS) on a purified MNdh-BMC-T^SE^ complex and BMC-T^SE^ alone, comparing the solvent accessible residues between the complex and isolated BMC-T^SE^. The samples were prepared anaerobically, and the low oxygen conditions were maintained by rapidly loading them into an airtight sample environment for microsecond X-ray exposure. Hydroxyl radicals are generated by radiolysis isotropically wherever water is present; thus, when water is near side chain residues, they yield covalently modified hydroxylated or carbonylated product, which generally requires an oxygen environment but is also known to form under a low oxygen environment^38^. We obtained near > 95% sequence coverage for BMC-T^SE^, and experiments in replicates exhibited consistent radiolytic labeling on identical residues. The hydroxyl radical reactivity rate constant was determined from the progressive extent of modification with an increase in the exposure time plots, which resembled the low oxygen environment for sample exposure (Fig. S5, Fig. S6, Table S2). Those residues identified with changes in their modification rate in the complex MNdh-BMC-T^SE^ vs. free BMC-T^SE^ are consequently likely to be involved in the conformational changes and interactions that mediate complex formation. The ratio of the site-specific rate constant of modification between isolated protein and complex provides the extent of solvent accessibility changes upon complex formation (Fig. 3). We divided the ratio of hydroxyl radical reactivities or changes in solvent accessibility into four categories. In the first category, Met23, Met100, and Met108, located in the hydrophobic core of individual subunits of BMC-T^SE^, showed the highest decrease (more than 40-fold) in solvent accessibility. Buried Met residues have been reported to show some degree of hydroxyl radical reactivity due to the dynamic nature of the core, which can momentarily expose this highly reactive residue to the bulk water, giving rise to its hydroxyl radical modification product^39^. However, here we observed almost complete protection of Met100 and Met108 upon formation of the complex. Since these residues project towards the same monomer core, between the beta-sheet and alpha helices of the BMC fold, this data indicates a nearly complete loss of dynamics upon MNdh binding, which could be due to a tightening of the interaction between subunits within the BMC-T^SE^ trimer upon binding MNdh. This is consistent with the disorder observed for the residues surrounding the pore in a crystal structure of the BMC-T^SE^ homolog PduT^40^.

**Figure 3.**
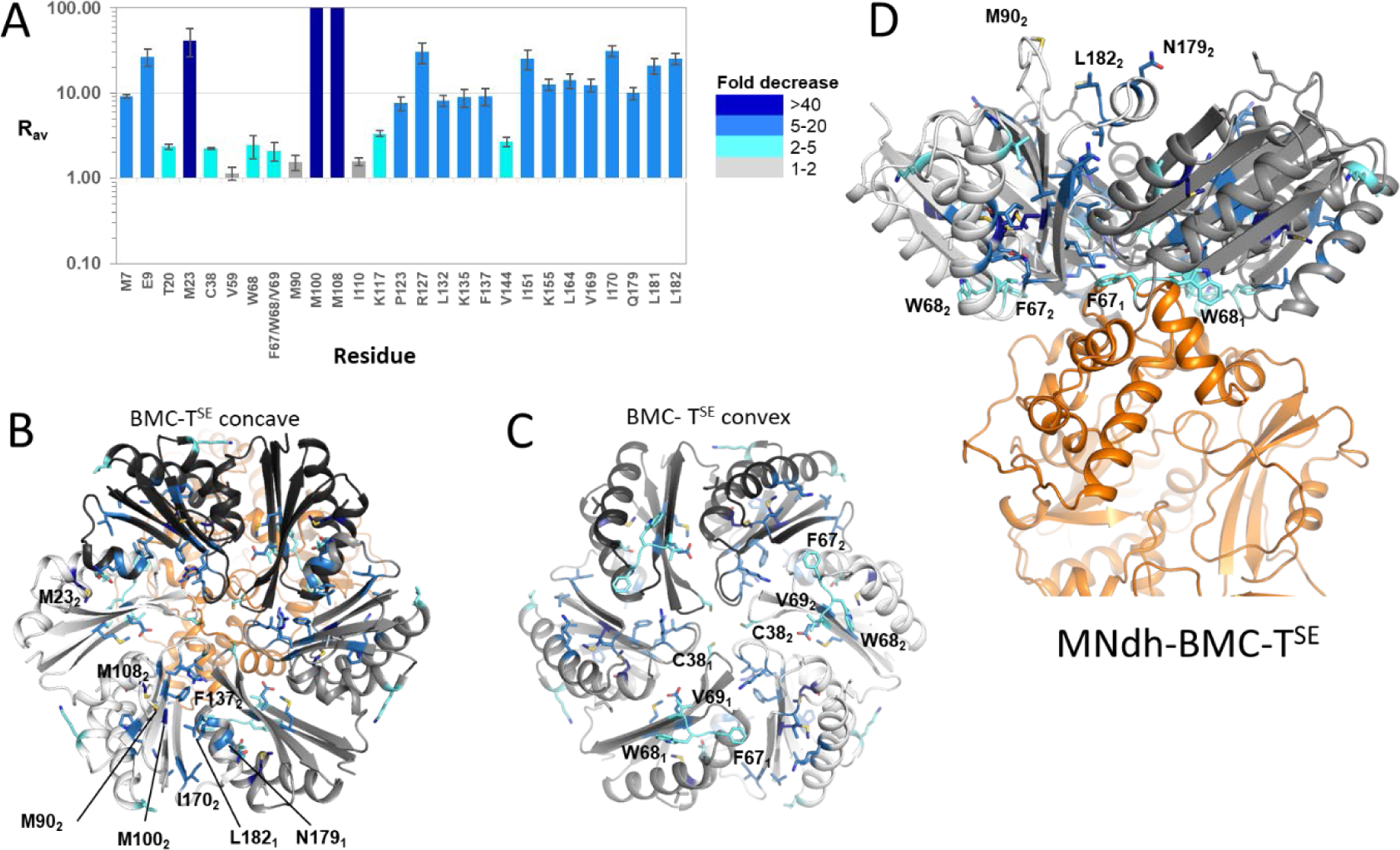
Structural characterization of the MNdh-BMC-TSE complex by X-ray footprinting. (**A**) Bar plot with color profile indicates fold change in the hydroxyl radical reactivity rate constant or solvent accessibility change between free BMC-TSE and MNdh-BMC-TSE complex. (**B**), (**C**), and (**D**) Residues of the BMC-TSE concave, convex, and side surface views are highlighted with the color profile in the bar plot and indicate fold change in solvent accessibility. Residue numbering follows the native protein sequence. Subscripts 1 and 2 highlight the asymmetric interaction of MNdh with BMC-TSE; the solvent accessibility of residues involved in the interaction represent an average over all subunits.

The vast majority of changes in solvent accessibility were in the second category, and included residues at the intermonomer interface or concave surface and intramonomer core: Met 7, Glu9, Pro123, Arg127, Leu132, Lys 135, Phe137, Ile151, Lys155, Leu164, Val169, Ile170, GLu179, Leu181, Leu 182. These residues showed a moderate decrease (∼ 10 fold) in solvent accessibility, indicating an overall loss of water accessibility upon formation of the MNdh-BMC-T^SE^ complex. A similar loss of hydroxyl radical reactivity in the concave surface was observed previously in the formation of BMC heterohexamers^41^. In the third category, residues Val59, Met90, and Ile110, which are not in direct contact with the subunit interface and MNdh, showed no change in solvent accessibility. In the fourth category, the residues Cys38, Phe67, Trp68, Val69, and three others (Thr20, Lys117, and Val144) showed only a ∼ 3-fold decrease in solvent accessibility (Fig. 3B). The AlphaFold2 model of the MNdh-BMC-T^SE^ complex indicates that only one out of three subunits of BMC-T^SE^ interact with MNdh near the Cys 38, Trp 68, Phe 67, and Val69 residues. XFMS reports on the protection pattern of the ensemble average under equilibrium conditions during the microsecond hydroxyl radical labeling reaction. Therefore, such nonspecific interactions will result in reduced sensitivity to differences in the reactivity rates and, thus, solvent accessibility changes. The approximate 1/3 solvent accessibility change at the hydrophobic cluster Phe 67-Trp 68-Val69 relative to the 10 fold change seen at the majority of residues supports the asymmetric interaction of MNdh with BMC-T^SE^ together with compaction of BMC-T^SE^ upon binding.

### Characterization of a purified MNdh-BMC-T^SE^ complex

We co-expressed MNdh and BMC-T^SE^ from *Clostridium botulinum* (a GRM1 type BMC) anaerobically with a histidine affinity tag on BMC-T^SE^. A mutation of lysine 25 (numbering of untagged protein sequence) to glutamine for BMC-T^SE^ was engineered to prevent self-assembly that was observed for a native complex. Affinity purification followed by size exclusion chromatography under anaerobic conditions allowed us to isolate a stable complex containing both proteins (Fig. 4A) that contained FMN and Fe-S clusters (Fig. 4B). The purified complex showed a rapid consumption of NADH measured by absorption when we incubate the complex with the substrate aerobically. This reaction is dependent on an available electron acceptor which appears to be oxygen, as the reaction only proceeds in air and anaerobically sealed solutions do not show appreciable NADH conversion unless exposed to the aerobic environment (Fig. 4D). Using a series of substrate concentrations at low enzyme concentrations we were able to measure initial rates at determine a k_cat_ of 20 s^-1^ and a K_m_ of 0.98mM for the reaction of NADH to NAD+ with oxygen as an electron acceptor (Fig. S7A). This is in line with similar rates for RnfC^42^; in this reaction we do not know, however, if the transfer of the electrons to oxygen occurs at the FMN site or at any of the 4Fe-4S clusters (or a combination of them). The reaction as observed for complex I is likely occurring at the site of the FMN ^43^. For the oxygen to access the FMN it is likely necessary for the NAD(H) site to be empty; consistent with this hypothesis is the observed inhibition of the oxygen reaction by the presence of excess substrate or product (see 0.15 μM MNdh/O_2_ trace in Fig. S7B that plateaus after 120 s). From our model (Fig. 2E), either the BMC-T^SE^ Fe-S cluster or the FMN seem the most likely candidates as electron donors due to their solvent exposure. The NADH to NAD+ conversion reaction in absence of any other substrate however provides support for our hypothesis that MNdh is not directly involved in CoenzymeA regeneration as previously proposed.

**Figure 4.**
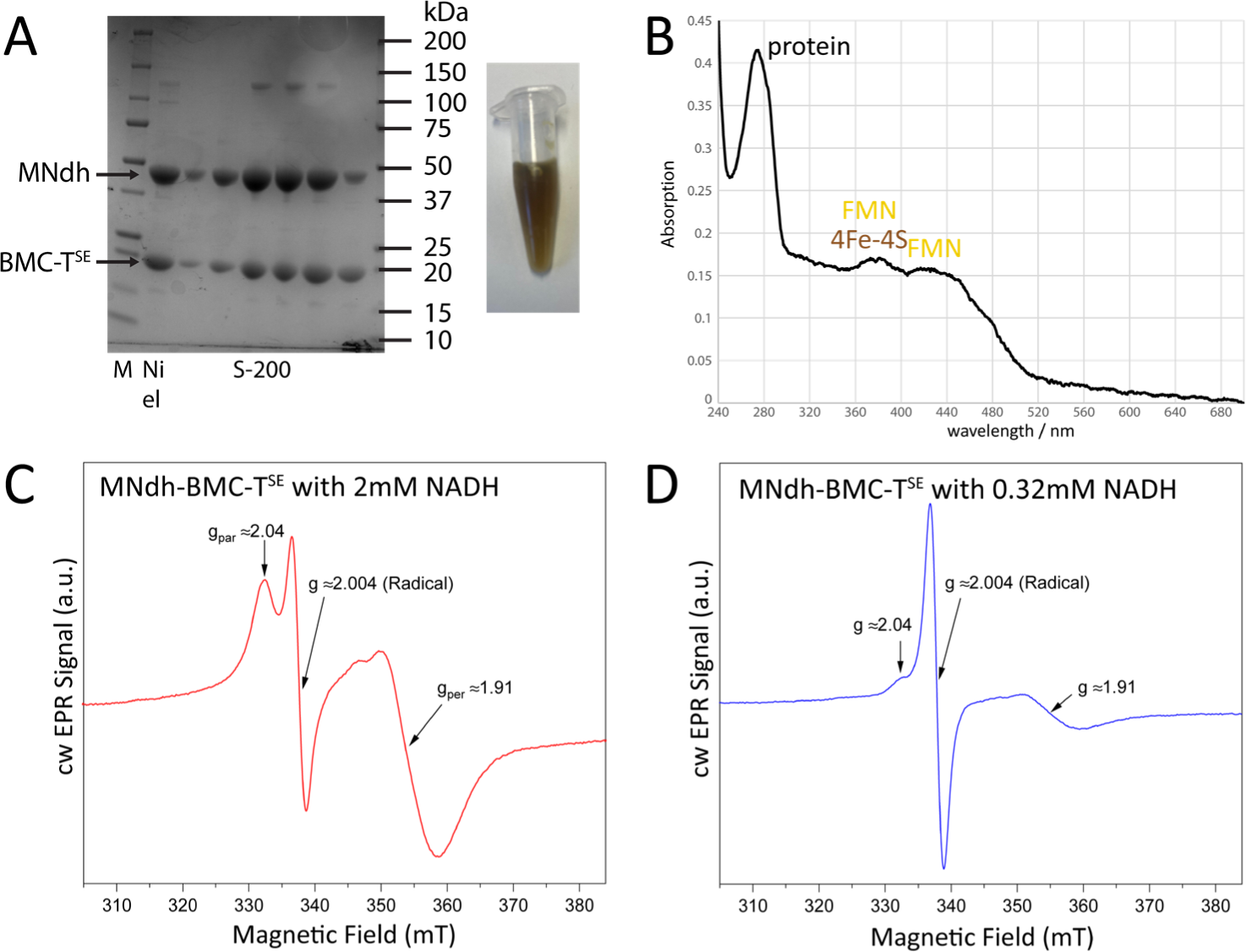
Characterization of the MNdh-BMC-TSE complex. (**A**) (left) SDS-PAGE of complex as observed after Ni affinity purification followed by S-200 size exclusion chromatography. (right) Color of purified complex indicates presence of Fe-S clusters and FMN cofactor. (**B**) UV-vis spectrum of the complex (5μM), showing absorption peaks typical for 4Fe-4S clusters and oxidized FMN. (**C**) Continuous wave (cw) X-band EPR Spectra of 0.32 mM MNdh-BMC-TSE complex reduced with 2 mM NADH (∼7x molar excess)., 3.2 mW microwave power, 1.2 mT modulation amplitude, 100 kHz modulation frequency, T= 10 K. (**D**) Same as (A) but MNdh-BMC-T^SE^ reduced with a stoichiometric amount of NADH (1:1). Note, that Fe-S Signal with g-factors gpar≈2.04 and gper≈1.91, is not saturated at this microwave power, while organic radical signal with gr≈2.004 is heavily saturated. Therefore, spin quantification for organic radical was done using EPR spectra recorded with significantly lower microwave power at higher temperature (see Fig. S8).

In order to verify our proposed hypothesis of electron transfer, we analyzed samples of the MNdh/BMC-T^SE^ complex sing continuous wave (cw) X-band Electron Paramagnetic Resonance (EPR) spectroscopy; samples were prepared under anaerobic conditions. Addition of 2mM (∼7x stoichiometric excess) of the reductant NADH to the 0.32 mM protein solution leads to the appearance of a strong, broad and fast relaxing signal characteristic for Fe-S clusters, with g-factors g_pa_r≈2.04 and gper≈1.91, and relatively small amount of organic radical with g ≈ 2.004 as shown in Fig. 4C. The obtained g-values for the Fe-S cluster are in excellent agreement with previous EPR experiments^17^ for BMC-T^SE^. Importantly, these values are different from those reported for Fe-S clusters in PduS alone^29^. Spin quantification of this Fe-S signal gave ≈0.17 mM, indicating that the reduction with NADH affected a substantial fraction of the protein. Concentration of the organic radical species was ≈0.007 mM which was determined by spin quantification of the spectra recorded at lower microwave power and higher temperature to avoid saturation behavior (Fig. S8). This concentration is roughly 45x lower than the protein concentration, showing that the radical is also formed substoichiometrically. The linewidth (≈2 mT) and g-value (2.004) of this radical signal are in agreement with a reduced FMN radical^44^.

Addition of a stoichiometric amount of NADH (0.32 mM) still leads to the appearance of Fe-S cluster signal with a concentration of ≈0.034 mM (Fig. 4D), which is ∼5 times less than in the 2mM NADH sample. The g-values of the Fe-S cluster signals in the two samples are similar, indicating that the same Fe-S cluster, i.e. the BMC-T^SE^ one, was reduced (Fig. 4D). A much stronger organic radical signal was observed in this sample and spin quantification gave 0.034 mM vs 0.007 mM spins (∼5 times stronger than the former sample). This implies that radical and Fe-S signals are formed in equal amount in about 1/10^th^ of the proteins. The1:1 ratio of radical to Fe-S signals in the sample reduced stoichiometrically with NADH can be explained by double reduction of FMN by NADH. Doubly reduced FMN (including its double protonated form FMNH_2_) is in a singlet spin state and thus cannot be observed by EPR. Further electron transfer to the Fe-S clusters of MNdh leads to the appearance of two EPR signals: one from singly reduced FMN radical and another one from singly reduced Fe-S cluster with stoichiometric ratio of 1:1.

To measure electrochemical properties of both FMN and the Fe-S clusters, we analyzed samples of the MNdh-BMC-T^SE^ complex using protein film square-wave voltammetry (SWV). Background-corrected SWVs of MNdh-BMC-T^SE^ reveal several overlapping redox peaks whose relative peak heights increase unevenly with increasing pulse frequency (Fig. 5A), thereby enabling deconvolution and identification of five distinct redox waves. This is consistent with three one-electron reductions associated with each Fe-S cluster, and two sequential one-electron reductions associated with FMN. Because FMN undergoes two sequential proton-coupled electron transfer steps, it can be modelled as a single adsorbed redox species and distinguished from the Fe-S clusters. Experimental protein film SWVs were fit to SWVs simulated using DigiElch and are in good agreement with a surface-confined model consisting of three one-electron reductions – standard reduction potentials of -0.147 V, -0.217 V, and -0.349 V (assigned here as the Fe-S clusters) – and two coupled one-electron reductions – standard reduction potentials of -0.284 V and - 0.459 V (assigned here as the quinone/semiquinone and semiquinone/hydroquinone forms of FMN). Furthermore, simulated SWV curves were fit with experiments across multiple frequencies using constant simulation parameters and were found to be in good agreement (R^2^ > 0.98 for all frequencies studied, Fig. 5B).

**Figure 5.**
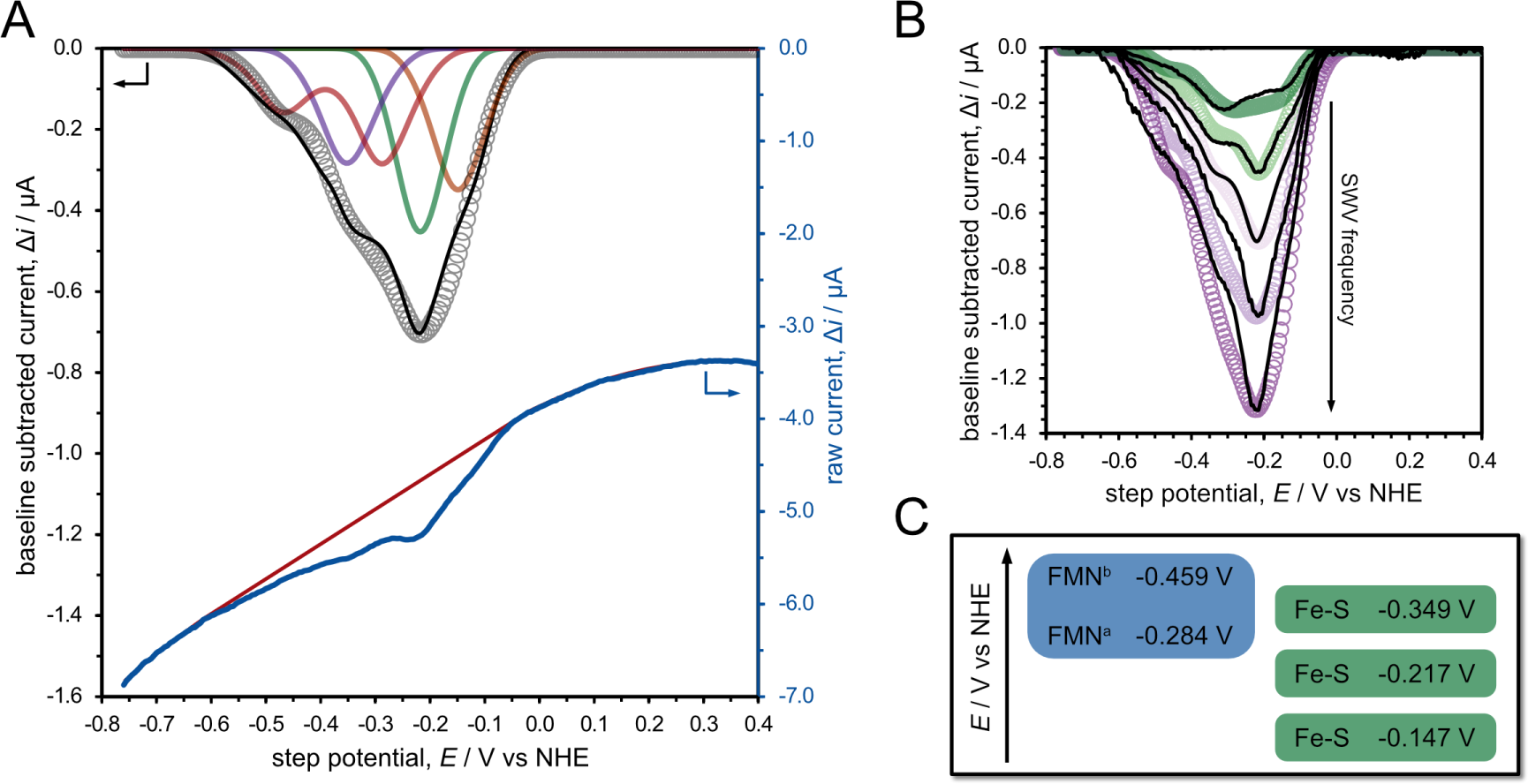
(**A**) Representative protein film SWV of MNdh-BMC-TSE complex (solid blue = raw current, solid red = background current, solid black = background-corrected current) and a SWV simulated using DigiElch (open grey circles) with its composite peaks (red, purple, green, and orange solid curves). Simulated SWVs are composed of four surface-confined species and five total redox waves: 3x reversible one-electron waves (orange, green and purple curves) corresponding to Fe-S clusters and 2x coupled one-electron EE waves (red curve) corresponding to the flavin. (**B**) Experimental (solid black curves) and simulated (open circles) protein film SWVs of MNdh-BMC-TSE complex at potential pulse frequencies of 5 Hz, 10 Hz, 20 Hz, 30 Hz, and 40 Hz. (**C**) Scheme illustrating the assigned redox potential for the Fe-S clusters and the first (FMNa = quinone/semiquinone) and second (FMNb = semiquinone/hydroquinone) redox potentials of the FMN cofactor. Electrochemical experiments were performed with a pulse height of 20 mV, step height of 5 mV, and frequency of 20 Hz (for **A**) using a HEPES-buffered saline solution under N2 at pH 7 and 25 °C.

Control SWV experiments performed using thermally denatured MNdh-BMC-T^SE^ reveal a single redox peak at -0.205 V consistent with free FMN^45,46^, suggesting that SWV peaks observed with intact MNdh-BMC-T^SE^ are not the result of fully denatured protein. Additionally, SWVs of partially denatured MNdh-BMC-T^SE^ (brief exposure to aerobic conditions to react the Fe-S clusters) reveal the loss of peaks at -0.147, -0.217, and -0.349 V while the peaks assigned to FMN (-0.284 V and -0.459 V) were unchanged (Fig. S9). Taken together, this data is consistent with the proposed assignment of redox peaks. Based on the assigned redox potentials, the first electron transfer from doubly reduced FMN to Fe-S is significantly more favorable than the second electron transfer from the semiquinone form. This relatively stable semiquinone may further explain the comparatively high concentration of organic radical observed by EPR after reaction with stoichiometric NADH. While formation of the stable semiquinone may not be relevant to the physiological function, shifts in the redox potentials when the complex is assembled into a shell and/or upon substrate binding could eliminate significant formation of the radical species.

## Discussion

The initial characterization of PduS, a MNdh homolog in PDU BMCs proposed a function as a regenerating enzyme for adenosylcobalamin / coenzyme B_12_ ^29^; however this cofactor is only required in the signature enzymes of the EUT and PDU BMCs. However, MNdh is widespread among BMCs and Coenzyme B_12_ does not seem to be a cofactor in any other characterized BMC types, nor is there evidence of B_12_ binding motifs in protein sequences associated with uncharacterized BMC types. The widespread occurrence of MNdh/BMC-T^SE^ across BMC types as well as across phyla (Fig. S1) strongly implies an alternative, more universal function. The initial proposed function was already contested at the time it was proposed^47^ and recently Burrichter *et al*.^13^ found a homolog of MNdh in a purified taurine degrading BMC and speculated that it could be involved in electron transfer via a BMC-T^SE^ protein.

There are five substantial pieces of evidence to support the function of MNdh as a housekeeping NAD+ regenerating BMC enzyme. (I) It is present in many types of BMCs that catabolize different substrates; these all contain aldehyde dehydrogenase; and many observations (361) in which the next step in the metabolosome paradigm catalyzed by alcohol dehydrogenase, with its required function of NAD recycling, is missing. (II) Homology to the function of structurally related enzymes RnfC and NuoF and high structural alignment over the whole sequence between RnfC and MNdh with little room for binding alternate cofactors such as cobalamin. (III) Distribution of MNdh and BMC-T^SE^ sequences on a phylogenetic tree is not correlated with BMC type, indicating a more general function unrelated to a specific substrate (Fig. S2). (IV) Strict co-occurrence of MNdh with BMC-T^SE^ (but not vice versa), highlighting the need for electrons to be passed to the outside if MNdh is present in the BMC. (V) Functional assays support the function of NAD+ regeneration *in vitro* without the need for additional substrates like Coenzyme B_12_.

There are likely a multitude of ways for cofactor regeneration in BMCs; at low throughput and with a high enough concentration gradient, BMCs might even work with cofactor exchange across the shell through some of the larger pores in BMC-T^S^ shell proteins. Many of the BMC loci that contain MNdh also contain an alcohol dehydrogenase (Fig. 1C, Table S1). Their functions are seemingly redundant, but complementary if dependent on redox condition. The MNdh reaction is likely faster since it only needs NADH to bind and not an additional substrate. Both the alcohol dehydrogenase as well as MNdh (PduS in PDU BMCs) have been analyzed in the *Salmonella* PDU system with genetic knockouts^6,31^. Despite the apparent necessity of an alcohol dehydrogenase for cofactor recycling inside the BMC, a PduQ (alcohol dehydrogenase) knockout was found to still grow on the PDU substrate 1,2-propanediol at 79% of the wildtype rate ^6^. Our proposed function for MNdh elegantly explains this activity that otherwise could only be explained with substantial amounts of NAD+/NADH transfer across the shell membrane. Likewise, the PduS knockout was found to also affect the growth rate^31^ but doesn’t abolish growth. This indicates that for *Salmonella* using PDU BMCs, under commonly used growth conditions, it is likely that both NAD+ regeneration systems are utilized. The redundancy attests to the importance of cofactor recycling for function of the BMC.

The observation of oxygen as a potential electron acceptor could provide a means to remove oxygen inside the BMC. Both MNdh and the alcohol dehydrogenase are sensitive to oxygen^6,29^. However, using oxygen as an electron acceptor results in reactive oxygen species but those might react non-specifically with proteins rather than with the sensitive Fe-S clusters. The strict co-occurrence of MNdh with BMC-T^SE^ (Table S1) suggests that they are both needed to fulfill the function of MNdh. This is consistent with our model of a linear electron transfer of two electrons from NADH to the FMN cofactor of MNdh and then through the two Fe-S clusters to the Fe-S cluster of BMC-T^SE^. We support this path and directionality with EPR measurements (Fig. 4CD) and protein voltammetry (Fig. 5AB). The distances between the Flavin and iron-sulfur clusters are short enough to allow this (Fig. 2E). Experimentally *in vitro* we have established oxygen as an electron acceptor (Fig. S7CD), however *in vivo* it is likely that a small redox protein, such as a ferredoxin or flavodoxin which could access the Fe-S cluster in the center of the concave BMC-T^SE^ pore are the final acceptors. There are a number of pfam02441 Flavoprotein type proteins in MNdh-encoding BMC operons and it is possible that some of those could act as electron acceptors. However those are also found in loci that do not contain MNdh, such as for the characterized flavodoxin from a PDU1C BMC locus^48^. The lack of a specific protein that co-occurs with MNdh/BMC-T^SE^ indicates that the interaction of BMC-T^SE^ with the acceptor is likely not very specific and different molecules are able to receive the electrons across the variety of species that MNdh/BMC-T^SE^ occur in. The lack of specificity is in line with what has been observed for soluble electron carriers in general^49^. It is however possible that oxygen can fill the role of electron acceptor *in vivo* as well. If this reaction occurs in proximity to the BMC that might be beneficial to protect other enzymes that are highly sensitive to oxygen such as the glycyl radical enzymes that undergo proteolytic cleavage when reacting with oxygen^50^. Oxygen as an electron acceptor has been observed for respiratory complex I in mitochondria and has major health implications due to the generation of superoxide, leading to cellular oxidative stress^51^.

There are a number of metabolosome BMC loci that contain neither of the regeneration systems, such as ARO, EUT3, MIC2/3, PVMlike, RMM/AAUM and SPU6^32^; how NAD+ regeneration is achieved for those cases has yet to be determined. Additional potentially redundant mechanisms for cofactor recycling underscores the importance of its maintenance for the function of these megadalton metabolic machines. There are also BMC types that do not have an AldDh such as the proposed Xau (Xanthine degrading BMC)^52^ where MNdh/BMC-T^SE^ are not found and there is no evidence of involvement of NADH for the reaction mechanism. Future work on specific BMC types should consider the mechanism for cofactor regeneration, which is just as important as a reaction pathway to explain the reactions occurring inside the BMC. The MNdh as a NAD+ regenerating enzyme improves the efficiency of the reactions by removing the need for an AlcDh that generates an alcohol side product for each turnover of the main substrate. The possibility of alternate electron acceptors, used under different physiological conditions, for this shell associated module suggests that it confers metabolic flexibility, which at the next step in the scale of organization is the hallmark metabolosome function. The new function of this enzyme in BMCs also established the BMC shell as a separating membrane for cofactor pools and conduit for electrons. This extends previous work on engineering metal centers into the pores of BMC shell proteins^53,54^ and sets the stage for connecting redox functions between the inside and outside of BMC shells.

## Materials and methods

### Cloning, protein expression, purification and activity assays

We have designed an *E. coli* codon optimized gene for MNdh and BMC-T^SE^ from the GRM1 operon of *Clostridium botulinum* E3 strain Alaska E43; see sequences in Table S3. The BMC-T^SE^ sequence contains an N-terminal his-tag and a mutation of the “antiparallel” lysine 25 (numbering of untagged protein sequence) to glutamine to prevent self-assembly. The open reading frames were cloned into a pACYCDuet vector using Gibson cloning.

The plasmid was transformed into ΔiscR BL21(DE3)^55^ *E. coli* cells for expression. 1.5 L LB broth medium with 0.5% glucose and buffered with 100mM MOPS/NaOH pH 7.4 was inoculated with 20ml of an overnight preculture and grown in a 2 L bottle in a shaking incubator at 120rpm and 37°C. At OD_600_ of 0.6-0.8 the culture was sparged with nitrogen and the medium was supplemented with L-cysteine at 2mM, sodium fumarate at 25mM, ferric ammonium citrate at 2mM and IPTG at 0.25mM final concentration. The flask was then sealed and grown overnight at 30°C. Cells were harvested by centrifugation at 7000 xg for 7 min and either processed immediately or frozen at -20°C until purification. For purification, the cell pellet from a 1.5L culture was resuspended in 20ml buffer A (20mM phosphate pH 7.0, 50mM NaCl, 20mM imidazole). 200µl 2mg/ml DNase I was added as well as ½ tablet of EDTA-free protease inhibitor (Roche). The cell suspension was lysed by two passages of a French Press at 25,000 psi, and the lysate was then cleared with a 20,000xg spin for 30 min. The soluble lysate was then transferred into a Coy anaerobic glove box under 2.5% hydrogen/nitrogen atmosphere and applied onto a 5ml HiTrap HP Ni-NTA resin, washed with 40ml buffer A and eluted with buffer B (20mM phosphate pH 7.0, 50mM NaCl, 300mM imidazole). The elution fraction was then concentrated to 500 µl using an Amicon concentrator with a 30kDa molecular weight cutoff (MWCO) and manually injected on a Superdex S-200 10/300 column equilibrated in 10mM PBS pH 7.0, 50mM NaCl and elution fractions were collected manually. Purification of BMC-T^SE^ alone was performed with the same protocol. Fractions were analyzed on SDS-PAGE and if necessary concentrated using and Amicon 30kDa MWCO concentrator. For enzymatic parameter determination assays, the protein was used at 4nM with NADH concentrations between 50μM to 750μM in a 96-well plate with 100μl volume per well and three replicates. The reaction was started by mixing protein and NADH with a large excess of aerobic buffer. Absorption at 340 nm was tracked using a Tecan Spark. The UV-Vis spectrum is an average of three traces of at 5 μM complex sample. Protein concentrations were determined using a BCA assay with a BSA standard curve.

### Bioinformatic analysis

The locus and protein sequence database from Sutter *et al.*^1^, available also as a web interface^32^, was used for all analysis. Protein sequences were of MNdh were aligned with ClustalW ^56^ trimming with trimAl 1.2rev59^57^ with parameters -gt 0.6 -cons 30 -w 3. The sequence alignment was then used to construct a maximum likelihood tree using RAxML-NG (v0.6.0)^58^. Trees were examined and visualized using Archaeopteryx (www.phylosoft.org/archaeopteryx).

### Structural modelling

The structure for the MNdh-BMC-T^SE^ complex was predicted using AlphaFold multimer^34,59^, as distributed in version 2.3.1 of AlphaFold. The prediction used the defaults with the full AlphaFold database and a template date of January 1^st^ 2022. AlphaFold predicted the BMC-T^SE^ trimer based on strong homology to existing solved structures for trimeric shell proteins. MNdh had fewer clear homologs in the PDB, but AlphaFold yields pLDDT scores above 70, and typically 90 or better (Fig. 2A). AlphaFold predicts that the MNdh monomer can be placed on either side of the trimer. Based on the anticipated localization for catalysis within the shell, only models where the MNdh was inside in the context of a BMC shell were considered realistic. Fe-S-clusters could be inserted manually in the AlphaFold apo structure with reasonable Cys-Fe distances. FMN and NAD+ were placed based on a structural alignment with the crystal structure of NuoF (pdb ID 6HL3), which contains both molecules. Protein structures were visualized with PyMOL (The PyMOL Molecular Graphics System, version 2.5.4 Schrödinger, LLC).

### X-ray radiolysis and mass spectrometry

Radiolysis of 8 µM BMC-T^SE^ and MNdh-BMC-T^SE^ samples were performed at the Advanced Light Source (ALS) beamline 3.3.1, which delivers a 3 – 12 keV broadband X-ray beam from a bending magnet source with a high flux density focused beam. We used an X-ray slit to produce a beam size at full-width-half-max of 640 μm (vertical) x 200 μm (horizontal) for sample exposure using a 200 μm ID capillary. Samples were loaded into a gas-tight lure lock Hamilton glass syringe and ejected through a 640 μm X-ray path using a 200 μm ID capillary at various speeds to achieve an exposure range of 400 to 1600 μs. Exposed sample was collected in tubes containing 60 mM methionine amide to immediately scavenge any secondary radical reaction products. The protein samples were cysteine alkylated and digested in trypsin and Glu-C using standard procedures^60^. LCMS was conducted on a Thermo Scientific Q Exactive Orbitrap Mass Spectrometer coupled to a Thermo Scientific UltiMate 3000 RSLCnano system. Digested samples were separated on a C18 column with a 15 minute acetonitrile/0.1% formic acid gradient as follows: ∼1–10% over 2.5 minutes, ∼10–40% over 12.5 minutes, and ∼40-90% over 1 minute. Peptides were analyzed in positive mode with data-dependent MS/MS using HCD fragmentation. Instrument control was via Xcalibur 4.1. XFMS peptide identification and analysis has been automated and enhanced by adopting the Byos® (Protein Metrics Inc) integrated software platform as previously described^61^. Biologic automatically extracts ion chromatograms and reports the quantification of modifications relative to the unmodified peptide based on the extracted ion chromatograms. A typical workflow starts with processing a high-exposure tandem MS (MS/MS) file in Byos for an MS/MS search against FASTA sequences and the localization of modification sites. The peptide level analysis and validation of assignments are carried out in Byologic and lead to the creation of in-silico peptides in the form of a CSV file using the MS/MS data. The in-silico peptides CSV is subsequently applied to full scan (MS1) data covering a series of exposure times, and the resulting quantified peptide modifications provide the basis for the residue-specific and peptide-level dose response. The abundance (peak area) of the identified unmodified and modified peptides at each irradiation time point area were measured from their respective extracted ion chromatogram of the mass spectrometry data collected in the precursor ion mode. The fraction unmodified for each peptide was calculated as the ratio of the integrated peak area of the unmodified peptide to the sum of integrated peak areas from the modified and unmodified peptides. The dose-response curves (fraction unmodified vs. X-ray exposure) were fitted to single exponential functions in Origin® Version 9.0 (OriginLabs). The rate constant, *k* (s^-1^), was used to measure the reactivity and solvent accessibility of sidechains towards hydroxyl radical-induced modification (Fig. S5, Fig. S6). The ratio of rate constants provided the relative change in the solvent accessibility between the free protein and the complex.

### EPR spectroscopy

Samples for EPR spectroscopy were prepared as above; the size exclusion peak fractions were concentrated to 0.2-0.3 mM (protein concentrations determined by BCA assay) using an Amicon spin concentrator and shipped/stored on ice until use. The EPR sample manipulation was done in a N_2_ glovebox to minimize damage due to oxygen exposure. The samples were filled into 4 mm o.d. thick-walled Suprasil X-band EPR tubes. Continuous wave (cw) X-band (9.5 GHz) EPR experiments were performed on a Bruker ELEXSYS II E500 EPR spectrometer (Bruker BioSpin, Ettlingen, Germany), equipped with a TE_102_ rectangular EPR resonator (Bruker ER 4102ST). The phase-sensitive detection with field modulation leads to first derivative type spectra, not the absorptive type spectra typically observed in other spectroscopic methods. All measurements were performed at cryogenic temperatures (10-30 K). A helium gas-flow cryostat (ICE Oxford, UK) and an ITC503 (Oxford Instruments, UK) were used for measurements at cryogenic temperatures. Data processing was performed using Xepr (Bruker BioSpin, Ettlingen) and Matlab (The MathWorks, Inc., Natick, Massachusetts, USA) software. Determination of the spin concentrations were done using the SpinCount application implemented in the Xepr software. Note that Fe-S signal at 10 K was not saturated at the microwave powers used in this study. The organic radical signal was strongly saturated at 10 K. For this reason the spin quantification of the organic radical was performed at T = 20 K, low microwave power, 6 μW, and small modulation amplitude, 0.4 mT (see Fig. S8).

### Protein Film Square Wave Voltammetry

Electrochemical experiments were performed using a BioLogic VMP potentiostat with a 3 mm glassy carbon working electrode, a Pt wire counter electrode, and Ag/AgCl reference electrode. Reference potentials were adjusted to the normal hydrogen electrode by adding 0.205 V. Working electrodes were cleaned and prepared following a protocol adapted from Plegaria *et al.*^54^. Electrodes were washed by sonication and subsequent rinsing with Milli-Q water. Protein film electrodes were prepared by drop casting 10 µL of 10mg/ml protein onto the electrode surface under a N_2_ environment and allowing them to dry over 15-30 minutes. Square wave voltammetry experiments were performed using 75 mM HEPES buffer with 150 mM NaCl at 25 °C under N_2_ atmosphere unless otherwise specified. Square wave voltammograms were performed from 0.2 V to -1.0 V vs Ag/AgCl with a step height of 5 mV, a pulse height of 20 mV, and pulse widths of 200, 100, 50, 33, and 25 ms (corresponding to 5, 10, 20, 30, and 40 Hz). For thermally denatured samples, protein was denatured at 95°C for 15 minutes before being drop cast and measured as above. Electrochemical data was processed (background subtraction and curve smoothing) using QSoas 3.2^62^.

Electrochemical simulations were performed using DigiElch 7.0 software. Protein films were modelled as surface-confined redox processes in a manner described by Mayall *et al.*^63^. Assuming that all species have the same surface concentration (i.e., cofactor stoichiometry is conserved on the electrode surface), the observed changes in relative peak heights with changes in pulse frequency can be attributed to differences in electron transfer rate constants between each cofactor and the electrode. A constant surface coverage was used for all simulations of Γ = 1.87 × 10^-11^ mol cm^-2^, while heterogeneous rate constants and redox potentials for each species were fit to the experimental SWV data.

## Acknowledgements

We thank Jasleen K. Bindra for help with the EPR measurements and Kalpana Singh for help with activity assays. This work was supported as part of the Center for Catalysis in Biomimetic Confinement, an Energy Frontier Research Center funded by the U.S. Department of Energy, Office of Science, Basic Energy Sciences under Award Number DE-SC0023395. The Advanced Light Source at Lawrence Berkeley National Laboratory is supported by the Director, Office of Science, Office of Basic Energy Sciences, US DOE under Contract DE-AC02-05CH11231. The EPR work is supported by the U. S. Department of Energy, Office of Science, Office of Basic Energy Sciences, Division of Chemical Sciences, Geosciences, and Biosciences, through Argonne National Laboratory under Contract No. DE-AC02-06CH11357. The Advanced Light Source and the Molecular Foundry are supported by the Office of Science of the U.S. Department of Energy (DOE) under contract DE-AC02-05CH11231. XFMS work was supported by NIH R01 GM126218 and NIH P30 GM124169.

## Author contributions

M.S. designed and performed experiments and interpreted results. C.A.K. designed and supervised the project and interpreted results. L.M.U., J.N and O.G.P. performed EPR spectroscopy and interpreted results. S.P., D.N.K., S.H. and C.Y.R. performed X-ray footprinting and interpreted results. B.F. performed molecular cloning, protein characterization and interpreted results. N.M.T. , M.A.T and D.P.H. performed protein voltammetry and interpreted results. J.V.V. performed protein structural modelling and interpreted results. M.S. and C.A.K. wrote the manuscript with help from all authors.

## Data and materials availability

All data are available in the main text or the supplementary materials.

