## Supplemental Figures and Tables for "Electrochemical cofactor recycling of bacterial microcompartments"

### Supplementary Materials

Fig. S1. Phylogenetic species tree with occurrences of MNdh.
Fig. S2. Phylogenetic tree of MNdh sequences and their distribution across bacterial phyla and BMC types.
Fig. S3. Alignment of RnfC structure to MNdh AlphaFold model.
Fig. S4. Sequence conservation mapped on AlphaFold2 models of BMC-$T^S$ from different types.
Fig. S5. Dose-response plots, part 1.
Fig. S6. Dose-response plots, part 2.
Fig. S7. Analysis of the NADH to NAD+ reaction.
Fig. S8. EPR spectra of FMN radical at low mw power.
Fig. S9. Square wave protein film voltammetry of MNdh-BMC-$T^{SE}$.
Table S1. Occurrence of MNdh across BMC types and co-occurrence with BMC-$T^{SE}$, AldDh and AlcDH.
Table S2. Rate constants of hydroxyl radical modification
Table S3. DNA and protein sequences

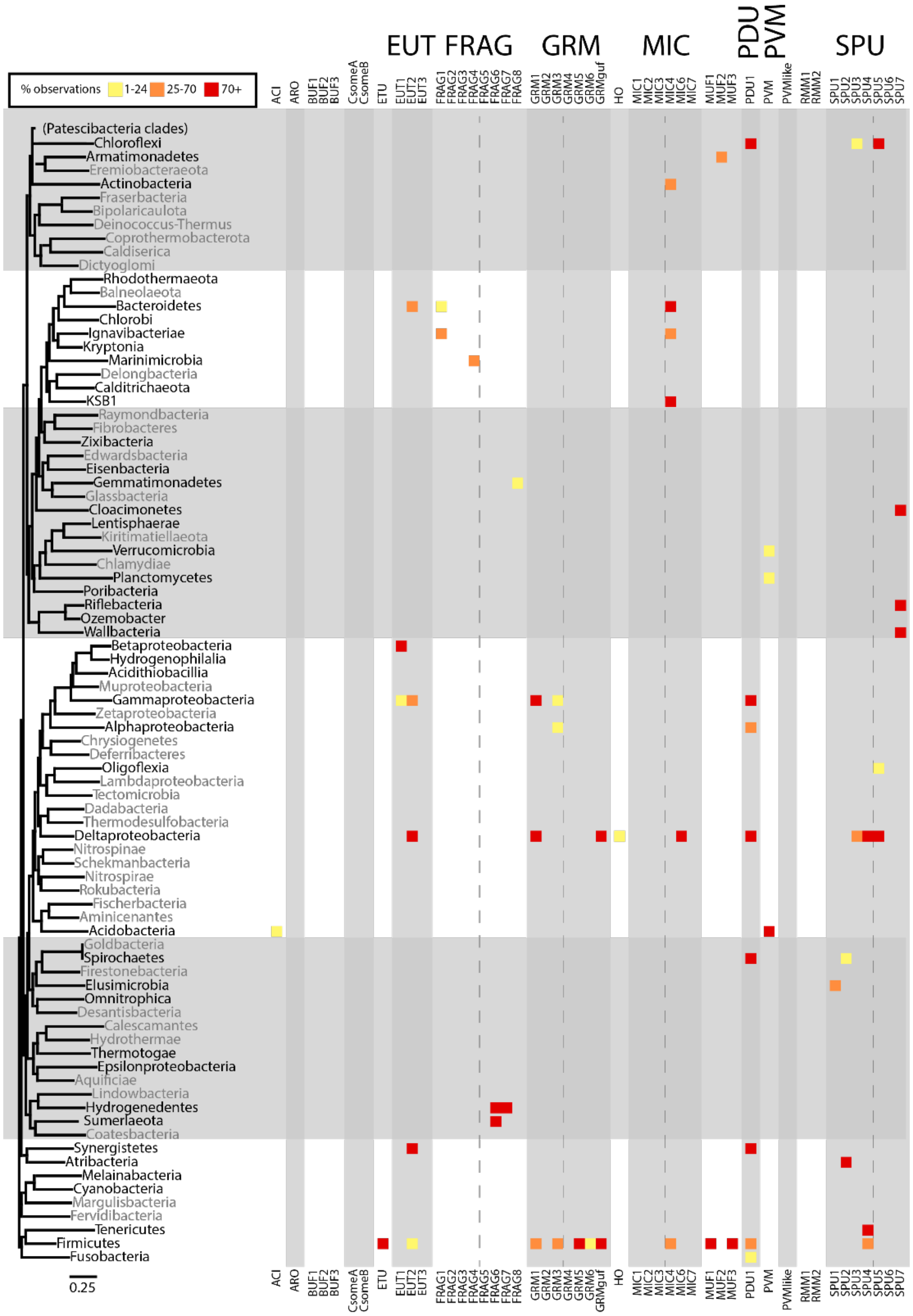

**Fig. S1. Phylogenetic species tree with occurrences of MNdh.** Prevalence of MNdh in BMC type, colored by percentage occurrence categories.

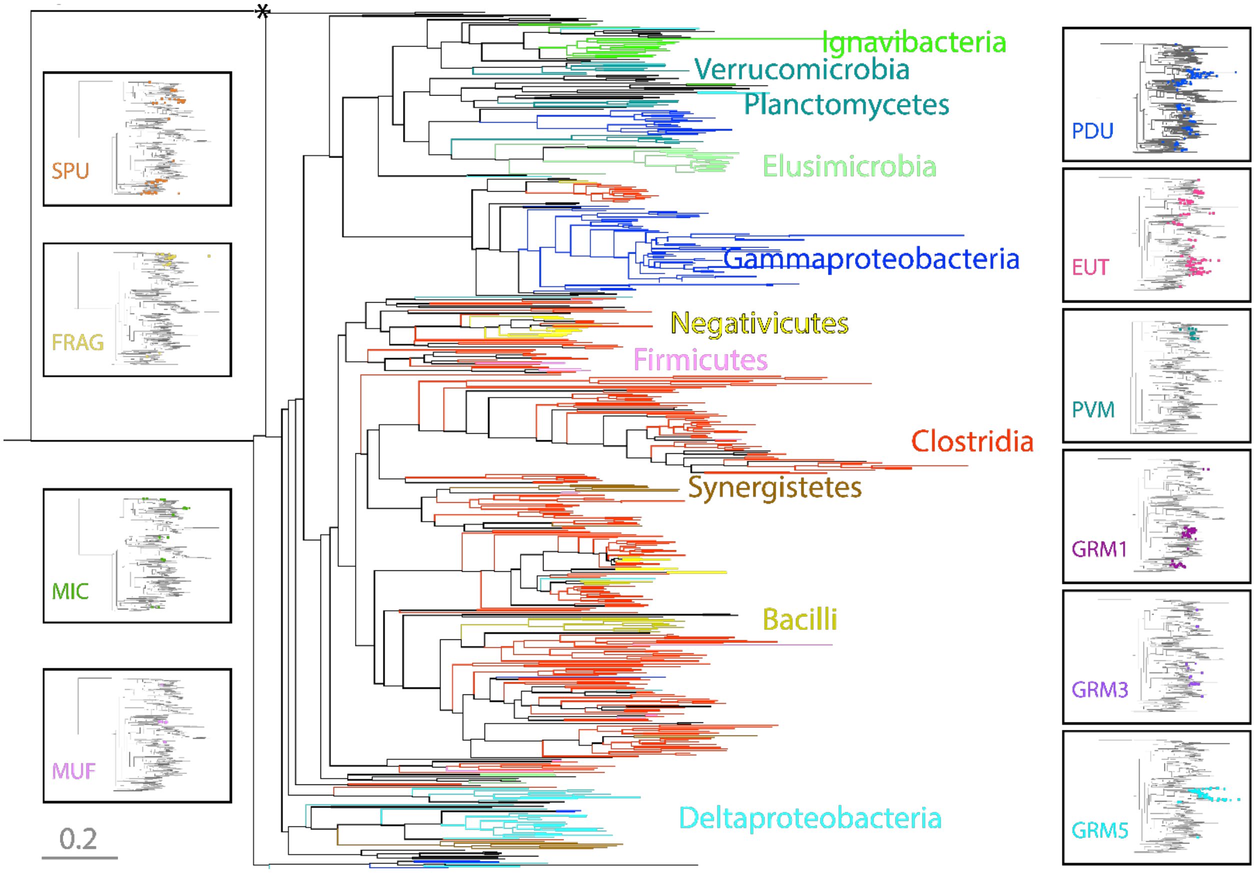


**Fig. S2. Phylogenetic tree of MNdh sequences and their distribution across bacterial phyla and BMC types.**

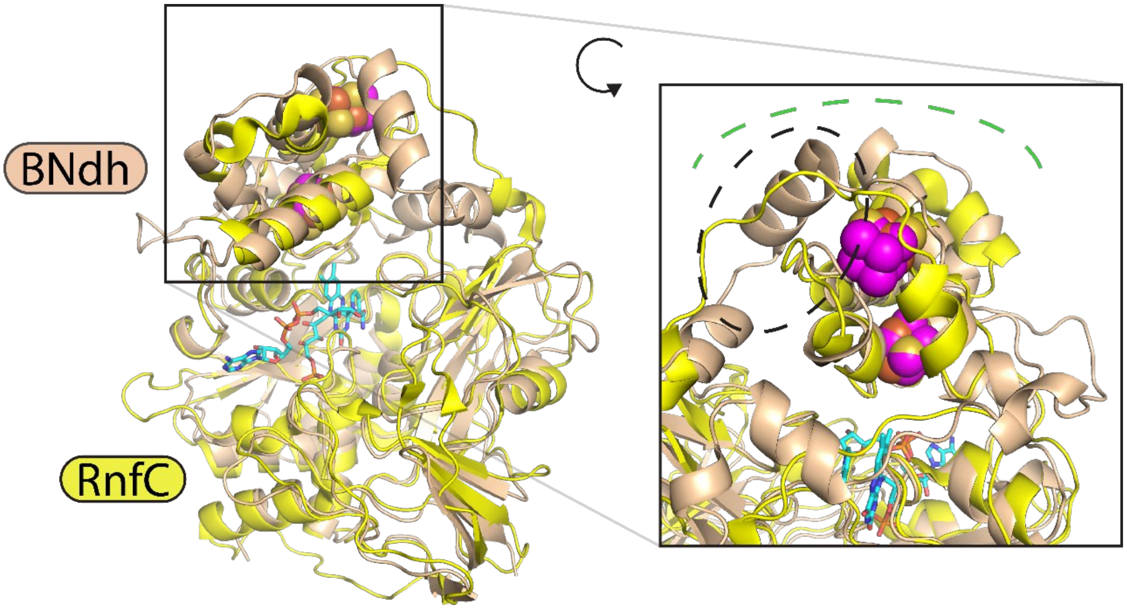


**Fig. S3. Alignment of RnfC structure to MNdh AlphaFold model.** Inset shows a closeup of the two 4Fe-4S clusters (RnfC clusters in magenta) and the region interacting with their respective partner proteins (green dash). The section with a dashed ellipse shows large difference between the RnfC structure and MNdh model.

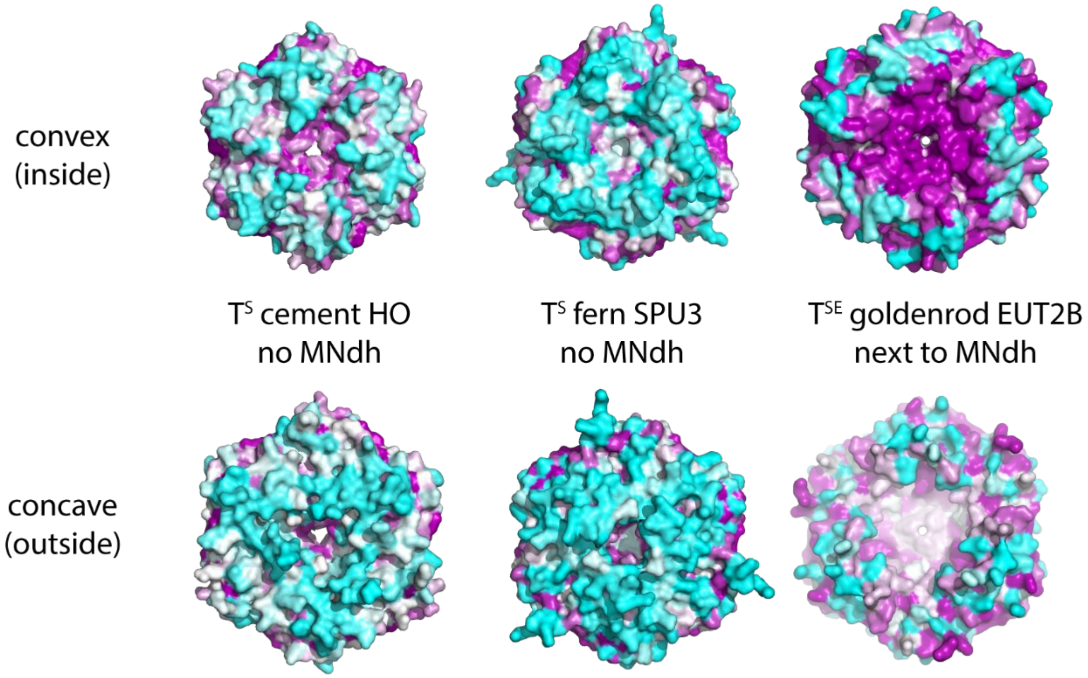


**Fig. S4. Sequence conservation mapped on AlphaFold2 models of BMC-$T^S$ from different types.** The BMC-$T^{SE}$ that co-occurs with MNdh shows much higher conservation around the pore region on the convex (interior) side. Sequence conservation is colored from low(cyan), medium (white) to high (magenta).

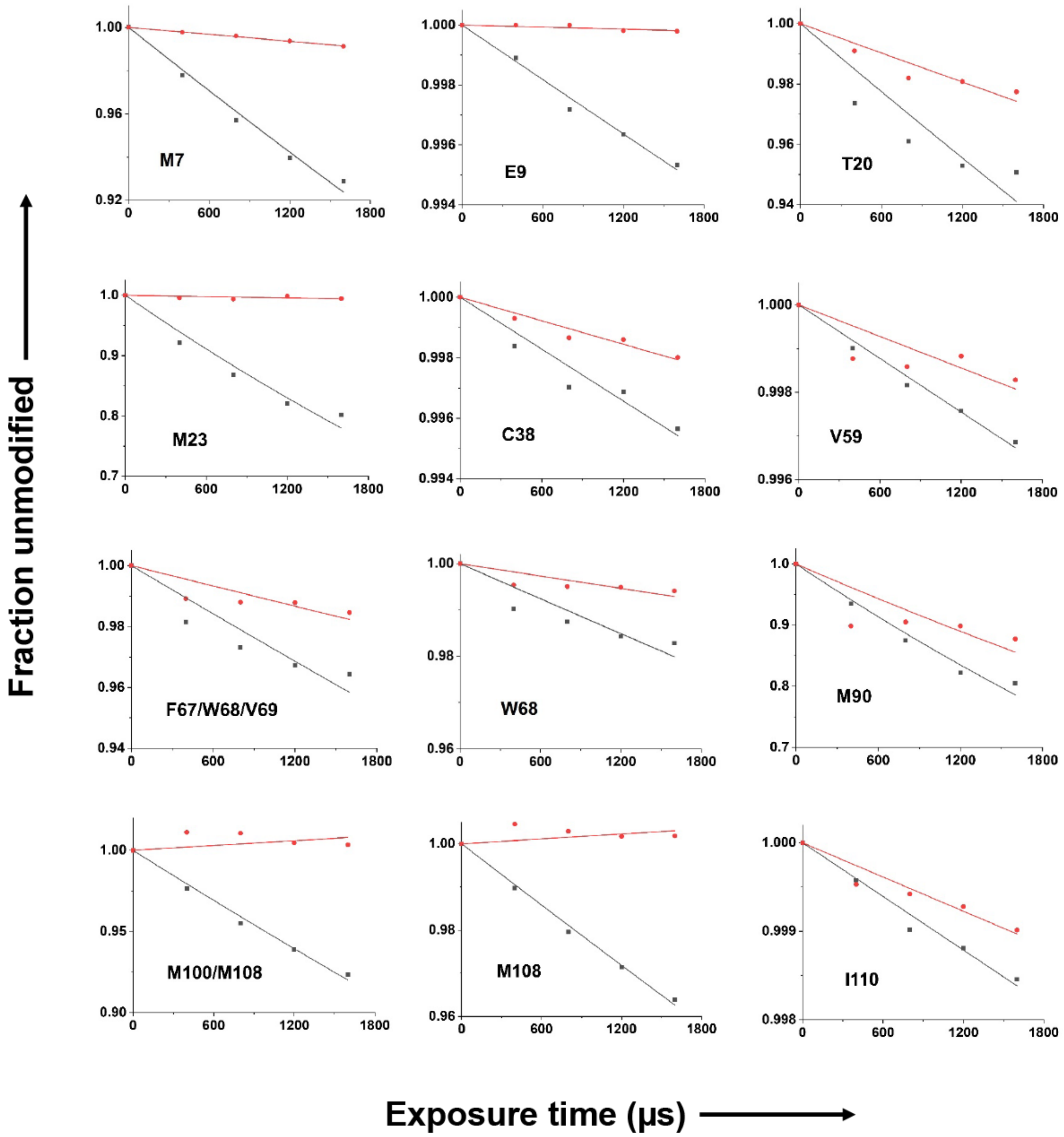


**Fig. S5. Dose-response plots, part 1.**

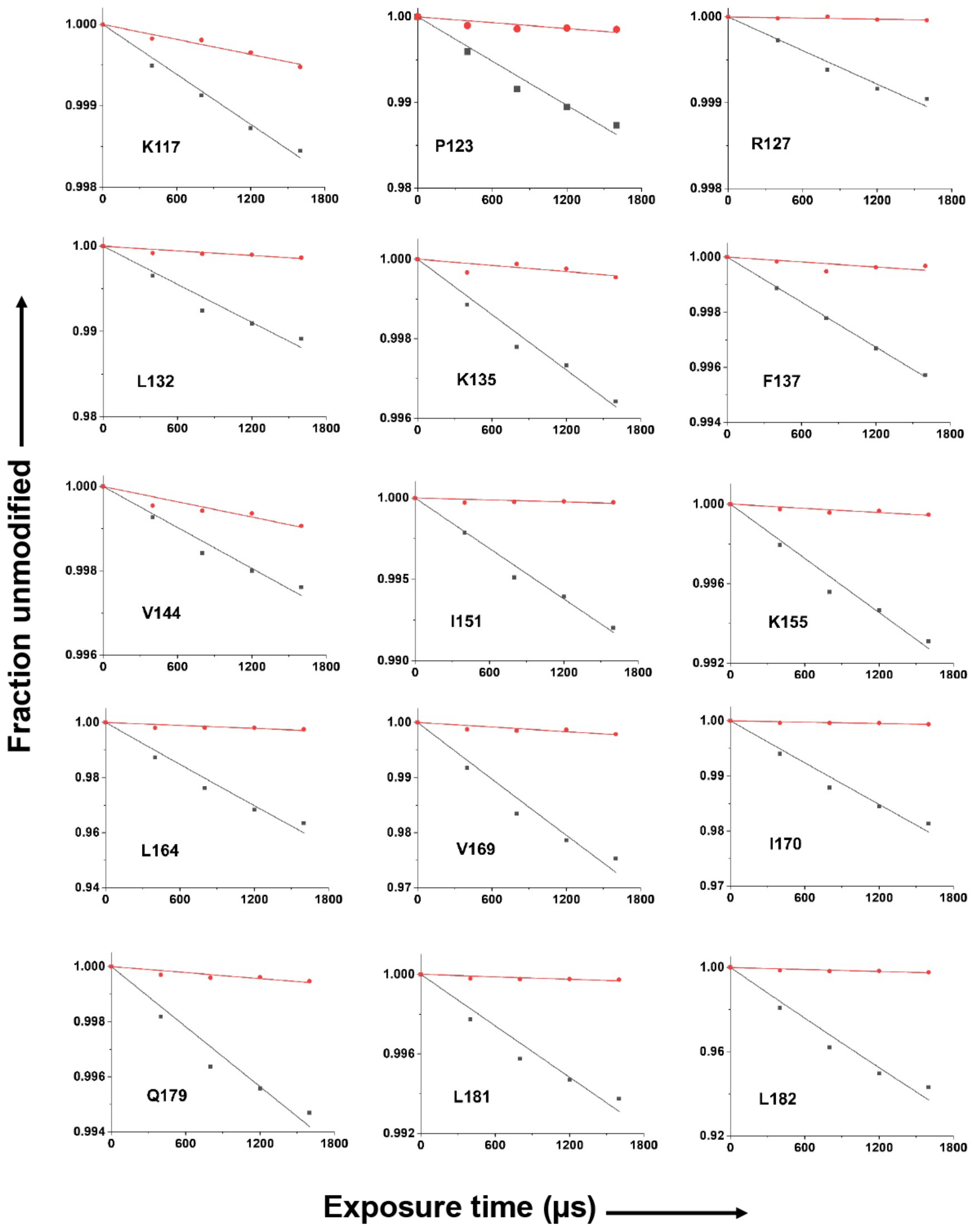


**Fig. S6. Dose-response plots, part 2.**

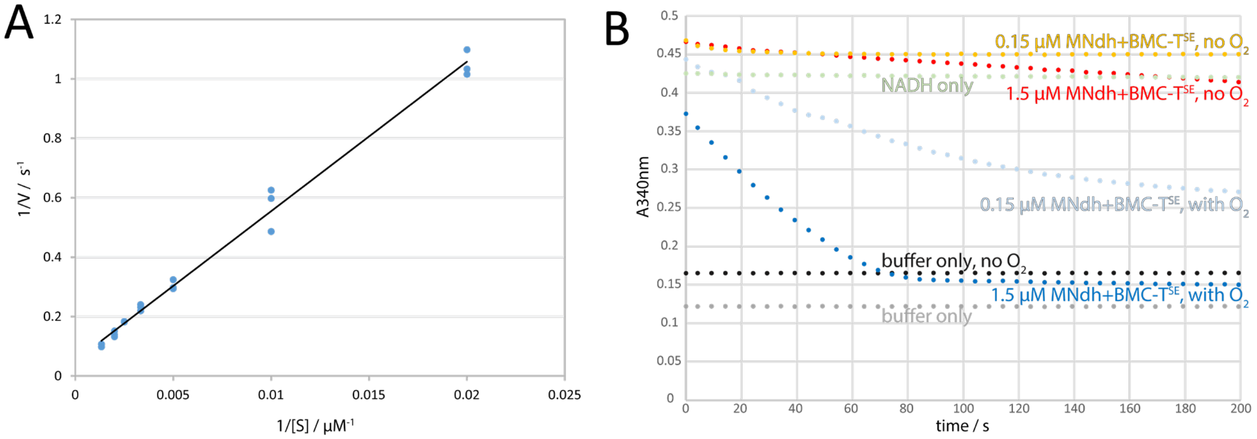


**Fig. S7. Analysis of the NADH to NAD+ reaction.** (A) Lineweaver-Burk plot of the NADH oxidation reaction with oxygen as electron acceptor. (**B**) Kinetic traces of the NADH oxidation reaction (0.4mM NADH) in the presence and absence of oxygen monitored by UV absorbance at 340nm.

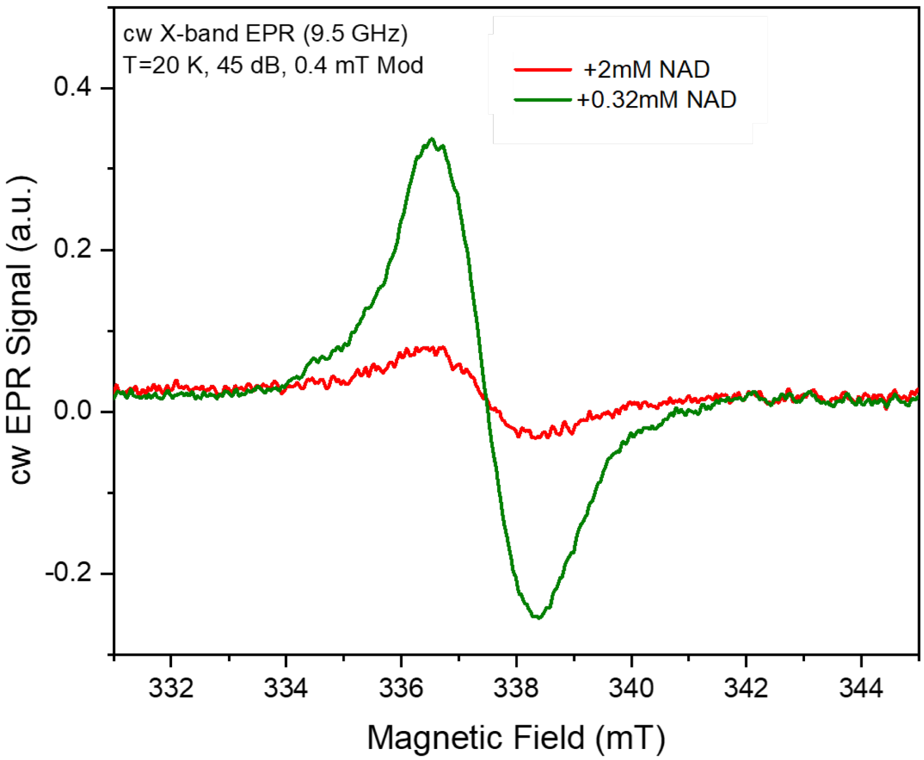


**Fig. S8. cw X-band EPR spectra of the organic radical recorded upon reduction of MNdh/BMC-T$^{SE}$ complex with NADH.** Spectra were recorded at low microwave power of 6 µW (45 dB attenuation of 200 mW), magnetic field modulation amplitude 0.4 mT, 100 kHz modulation frequency, T = 20 K. Red – reduction with 7 times molar excess of NADH (2 mM). Green - reduction with stoichiometric amount of NADH (0.32 mM).

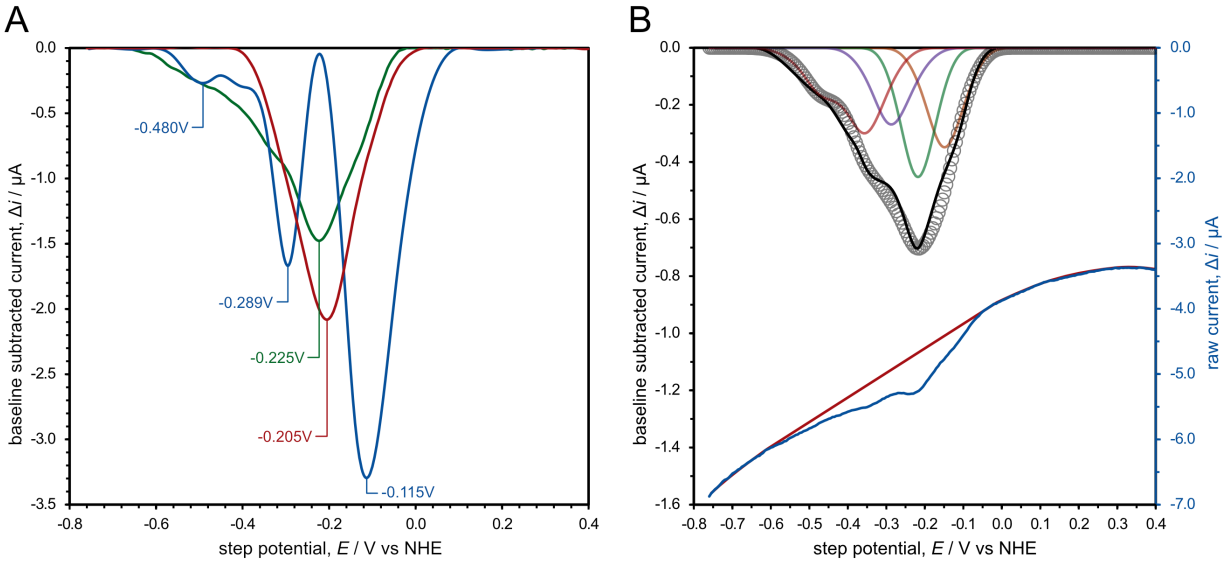


**Fig. S9. Square wave protein film voltammetry of MNdh-BMC-T$^{SE}$**. (**A**) Representative background corrected protein film SWVs of MNdh-BMC-T$^{SE}$ complex maintained under anaerobic conditions (green), exposed to air (blue), or thermally denatured (red). For MNdh-BMC-T$^{SE}$ complex exposed to air, the two redox peaks assigned to FMN (-0.480 V and -0.289 V) are largely unaffected, while the peaks assigned to Fe-S clusters (-0.354 V, -0.220 V, and -0.149 V) decrease concomitant with a broad asymmetric redox wave with a peak at -0.115 V (consistent with partial denaturation of the Fe-S cofactors upon exposure to ambient oxygen). Protein film SWV of thermally denatured MNdh-BMC-T$^{SE}$ complex reveals a single redox wave (-0.205 V) consistent with free FMN. (**B**) Representative protein film SWV of MNdh-BMC-T$^{SE}$ complex (solid blue = raw current, solid black = background-corrected current) and a simulated SWV (open grey circles) with its composite peaks (red, purple, green, and orange solid curves). The simulated SWV is composed of four surface-confined species and five total redox waves (3x reversible one-electron waves representing the Fe-S clusters and 1x coupled two-electron EE wave representing the flavin). Unlike the simulation presented in Fig. 5, the coupled redox peaks shown here are the two most reducing peaks (-0.35 V and -0.46 V). This model fits the experimental data nearly as well as that shown in Fig. 5 ($R^2$ = 0.983 vs $R^2$ = 0.984); however, it is not consistent with the control experiments nor the EPR data. No other parameters were found to accurately fit experimental SWVs at both high and low frequencies. Electrochemical experiments were performed with a pulse height of 20 mV, step height of 5 mV, and frequency of either (**A**) 50 Hz or (**B**) 20 Hz using a HEPES-buffered saline solution under $N_2$ at pH 7 and 25 °C.

| BMC type | total number of loci | percentage of loci with MNdh | number of loci with MNdh | loci with MNdh and BMC-T$^{SE}$ | loci with MNdh and AldDh | loci with MNdh and AlcDh |
|---|---|---|---|---|---|---|
| ACI | 125 | 6% | 8 | 7 | 7 | 0 |
| ETU | 3 | 100% | 3 | 3 | 2 | 3 |
| EUT1 | 1376 | 7% | 90 | 85 | 89 | 82 |
| EUT2 | 815 | 18% | 145 | 144 | 142 | 50 |
| FRAG | 117 | 24% | 28 | 27 | 5 | 1 |
| GRM1 | 318 | 31% | 97 | 94 | 95 | 43 |
| GRM3 | 200 | 28% | 56 | 55 | 56 | 54 |
| GRM5 | 245 | 98% | 240 | 236 | 222 | 233 |
| GRM6 | 19 | 11% | 2 | 2 | 2 | 2 |
| GRMguf | 18 | 100% | 18 | 18 | 18 | 4 |
| HO | 41 | 7% | 3 | 3 | 3 | 0 |
| MIC4 | 15 | 87% | 13 | 13 | 11 | 0 |
| MIC6 | 3 | 100% | 3 | 3 | 3 | 0 |
| MIC7 | 8 | 50% | 4 | 4 | 4 | 0 |
| MUF1 | 7 | 100% | 7 | 7 | 7 | 7 |
| MUF2 | 19 | 5% | 1 | 1 | 1 | 0 |
| MUF3 | 9 | 89% | 8 | 8 | 8 | 0 |
| PDU | 1649 | 81% | 1338 | 1330 | 1331 | 1278 |
| PVM | 285 | 8% | 22 | 22 | 21 | 0 |
| SPU1 | 62 | 65% | 40 | 40 | 36 | 0 |
| SPU2 | 53 | 25% | 13 | 12 | 12 | 1 |
| SPU3 | 70 | 6% | 4 | 4 | 4 | 0 |
| SPU4 | 33 | 42% | 14 | 14 | 11 | 7 |
| SPU5 | 8 | 88% | 7 | 7 | 7 | 0 |
| SPU7 | 8 | 100% | 8 | 8 | 8 | 0 |

**Table S1. Occurrence of MNdh across BMC types and co-occurrence with BMC-T$^{SE}$, AldDh and AlcDH.** Marked in orange are high prevalence of MNdh in BMC types, marked in light blue a high co-occurrence of MNdh and alcohol dehydrogenase.

| Sequence of the peptide fragment[a] | Site of modi-fication[b] | Type of modification(Da)[c] | Hydroxyl radical reactivity rate $k$ ($s^{-1}$)[d] | | R[e] | $R_{max}$[e] | $R_{min}$[e] | $R_{av}$[e] |
|---|---|---|---|---|---|---|---|---|
| | | | BMC-T[SE] | MNdh-BMC-T[SE] | | | | |
| [4]AIGMVEFISISR[15] | M7 | +16 | 49.54 ± 1.82 | 5.39 ± 0.09 | 9.20 | 9.70 | 8.71 | 9.20 ± 0.40 |
| | E9 | -30 | 3.03 ± 0.10 | 0.11 ± 0.03 | 26.78 | 35.82 | 21.09 | 28.46 ± 6.07 |
| [16]GIYATDQMLQTSD VEIVTANSVCPGK[41] | T20 | +16 | 53.31 ± 4.90 | 22.78 ± 0.03 | 2.34 | 2.56 | 2.12 | 2.34 ± 0.18 |
| | M23 | +16 | 155.35 ± 9.36 | 3.73 ± 1.25 | 41.65 | 66.04 | 29.40 | 47.72 ± 15.23 |
| [31]IVTANSVCPGKYIAI VNGDVAAVKE[55] | C38 | +48 | 3.78 ± 0.08 | 1.69 ± 0.02 | 2.24 | 2.31 | 2.16 | 2.24 ± 0.06 |
| [55]ESVSVGEK[62] | V59 | +14 | 2.34 ± 0.05 | 2.03 ± 0.37 | 1.15 | 1.44 | 0.95 | 1.19 ± 0.20 |
| [63]IAGEFWVDSIIIPN[76] | F67/W68 /V69 | | 36.64 ± 3.60 | 17.6 ± 3.43 | 2.08 | 2.84 | 1.57 | 2.21 ± 0.73 |
| | W68 | +32 | 17.58 ± 2.48 | 7.25 ± 1.51 | 2.42 | 3.49 | 1.72 | 2.61 ± 0.52 |
| [71]SIIIPNVNPLVFPAIT GATMPD[92] | M90 | +16 | 150.57 ± 7.28 | 97.38 ± 18.24 | 1.55 | 1.99 | 1.24 | 1.62 ± 0.31 |
| [98]GIMESFSLATMVIA ADAILK[117] | M100 /M108 | +16 | 52.08 ± 1.47 | 0.001[f] | ∞[f] | -[f] | - | - |
| [94]SLATMVIAADAILK[117] | M108 | +16 | 23.81± 0.51 | 0.001 | ∞ | - | - | - |
| | I110 | +16 | 1.01± 0.04 | 0.64 ± 0.05 | 1.57 | 1.78 | 1.39 | 1.58 ± 0.16 |
| | K117 | +16 | 1.02 ± 0.03 | 0.31 ± 0.02 | 3.34 | 3.68 | 3.04 | 3.36 ± 0.26 |
| [118]AANLEPLDLR[127] | P123 | +16 | 8.65 ± 0.46 | 1.14 ± 0.19 | 7.60 | 9.59 | 6.17 | 7.88 ± 1.40 |
| | R127 | +16 | 0.65 ± 0.03 | 0.02 ± 0.01 | 30.30 | 42.99 | 22.86 | 32.92 ± 8.31 |
| [128]LGTGLGGK[135] | L132 | +16 | 7.48 ± 0.46 | 0.92 ± 0.11 | 8.14 | 9.86 | 6.79 | 8.33 ± 1.26 |
| | K135 | +16 | 2.32 ± 0.10 | 0.26 ± 0.06 | 8.91 | 12.04 | 6.94 | 9.49 ± 2.10 |
| [136]SFFTF[140] | F137 | +16 | 2.73 ± 0.02 | 0.30 ± 0.07 | 9.14 | 12.27 | 7.26 | 9.76 ± 2.07 |
| [141]TGDVAAVEAGIDA GK[155] | V144 | +16 | 1.62 ± 0.08 | 0.60 ± 0.06 | 2.68 | 3.11 | 2.33 | 2.72 ± 0.32 |
| | I151 | +16 | 5.18 ± 0.20 | 0.20 ± 0.05 | 25.49 | 35.38 | 19.60 | 27.49 ± 6.51 |
| | K155 | +16 | 4.55 ± 0.21 | 0.36 ± 0.05 | 12.79 | 15.43 | 10.78 | 13.11 ± 1.91 |
| [162]GLLVNAEVIPSPS DR[176] | L164 | +16 | 25.54 ± 1.37 | 1.81 ± 0.32 | 14.12 | 18.11 | 11.34 | 14.72 ± 2.78 |
| | V169 | +16 | 17.23 ± 0.96 | 1.38 ± 0.20 | 12.46 | 15.36 | 10.28 | 12.82 ± 2.09 |
| | I170 | +16 | 12.73 ± 0.63 | 0.41 ± 0.05 | 31.36 | 37.94 | 26.32 | 32.13 ± 4.76 |
| [177]LLQSLL[182] | Q179 | +16 | 3.64 ± 0.22 | 0.36 ± .05 | 10.02 | 12.18 | 8.35 | 10.27 ±1.57 |
| | L181 | +14 | 4.32 ± 0.26 | 0.20 ± 0.04 | 21.18 | 27.30 | 16.90 | 22.10 ± 4.27 |
| | L182 | +16 | 40.59 ± 2.29 | 1.59 ± 0.21 | 25.45 | 31.00 | 21.21 | 26.10 ± 4.01 |

**Table S2. Rate constants of hydroxyl radical modification**

[a] sequences of tryptic fragments used for identification of quantification of modification sites

[b] modified residues, which were identified and confirmed by LCMS/MS

[c] type of side chain modification include hydroxylation, carbonylation, and decarboxylation, which resulted in a mass shift of +16, +14 and -30 Da

[d] hydroxyl radical rate constants were estimated by employing a first-order exponential fit of the dose-response plot of overall peptide modification as described in experimental procedures and Fig. 3. The modified peptide fragments were eluted as single or multiple peaks. The modified peak areas were extracted individually but summed together to calculate the total modification of the respective peptide.

[e] ratio of hydroxyl radical reactivity obtained by dividing hydroxyl radical reactivity rate of free by that of for the complex. The ratio represents fold decrease (>1) or increase (<1) in the solvent accessibility of the modified residues. $R_{max}$ and $R_{min}$ are the maximum and minimum values of the ratio based on the error of the hydroxyl racial reactivity constant. $R_{av}$ is the average value of R, $R_{max,}$ and $R_{min.}$

[f] value to show that the hydroxyl radical reactivity is near "zero", therefore the ratio is infinity, and we could not measure the $R_{max}$ and $R_{min}$

| Name | DNA/protein | Sequence |
|---|---|---|
| Cbot_MNdh | DNA | ATGTCCTTGTTGGATATGGTAAAAGACGCTGGAGTTATTGGTGCTGGGGGAGCAGGATTCCCCACACATGC<br>CAAGTTGGCATCCAAAGCGGAATACACCCTGTTGAACGGCGCCGAGTGTGAACCGCTTCTTCGTGTGGACC<br>AACAATTAATGGAATTGTTCCCGGACGAGATTATCAAGGGATTCGAGACTGCCCGTTCGATTGTCGACGCA<br>AACAAGGGGATTATCGGCATTAAAGGAAAGCATAAAGAGGTAATTTCGATTCTTCGTAATCGTATCAACGA<br>ATTACATGTTGGTGATCTTATCGAGGTTAAGGAGTTACCCGATATCTACCCTGCTGGTGACGAGCAAGTGT<br>TGGTGTATGAGTTAACTGGCCGTGTCGTCCCAGAGGCAGGCATCCCTATCCAGGTGGGATGCGTGGTAGTT<br>AACTCAGAAACCGCTTTAAATATCTACAATGCATCAATGGGCAAATCGGTTACAGAAAAGTATATTACAAT<br>CGCAGGAGATATTCCTAACCGCAAGACCGTAAAAGTACCGGTAGGGATTCCCATTATTGACGTGTTAAAGT<br>TGAGCGGGATTGAAAATTTCGATGACTATTCTGTTATTGACGGTGGCCCTATGATGGGCCCAGTCATGAAG<br>AACCTGGATGGGTATGTGACGAAGAAAAATAAGGGCTTTGTAATTCTTAAAAATGACCATCCTTTGATTCG<br>CAAGAAGAGCGTAACCATGAATCAAGCTAAAAAAATTAATAAGAGCTCCTGCGAACAGTGCCGTATGTGTA<br>CGGACATGTGTCCCCGCTTCTTACTTGGACACAACACCCAACCTCATAAGATGATGCGCGCCATGTCCTAT<br>AACTTGGACAACATTGAAGGGCAAAAGATTGCACAGCTGTGCTGTCAATGTAATTTATGCGAATTGTTTTC<br>CTGTCCCGCCGGTCTGTACCCGAAGTCTTCTAACTTATATTTCAAACAAAAGCTTGCTGAGCAGAACATTC<br>GCTATAAACCTGTACAGGACAAATTCGAGGCTCGTCAGGCTCGTGAATACCGCCTTGTCCCTAGCAAACGT<br>TTAATTGCACGTCTGGGGTTACGTGACTTCGATAAACCGGCCCCTATGATTGACGGACTTGTTGACTCCGA<br>GATTGTCTATATCGCCATGCAGCAGCACGTTGGCGCACCATCTATTGCGTGCGTGCACCTTGGACAACATG<br>TTGAGAATGGTGAGGTTATCGGCAAAATTCCCGAGGGATCTTTGGGCGCCAGTATCCATGCATCGATCAGT<br>GGTACTGTCATCAAAAACGAAGATGGATTTGTCGCTATCCGTCGCGACTAA |
| Cbot_MNdh | protein | MSLLDMVKDAGVIGAGGAGFPTHAKLASKAEYTLLNGAECEPLLRVDQQLMELFPDEIIKGFETARSIVDA<br>NKGIIGIKGKHKEVISILRNRINELHVGDLIEVKELPDIYPAGDEQVLVYELTGRVVPEAGIPIQVGCVVV<br>NSETALNIYNASMGKSVTEKYITIAGDIPNRKTVKVPVGIPIIDVLKLSGIENFDDYSVIDGGPMMGPVMK<br>NLDGYVTKKNKGFVILKNDHPLIRKKSVTMNQAKKINKSSCEQCRMCTDMCPRFLLGHNTQPHKMMRAMSY<br>NLDNIEGQKIAQLCCQCNLCELFSCPAGLYPKSSNLYFKQKLAEQNIRYKPVQDKFEARQAREYRLVPSKR<br>LIARLGLRDFDKPAPMIDGLVDSEIVYIAMQQHVGAPSIACVHLGQHVENGEVIGKIPEGSLGASIHASIS<br>GTVIKNEDGFVAIRRD* |
| Cbot_hisBMC<br>TSE_K25Q | DNA | ATGGGAAGCCACCATCACCATCATCATGGAAGCGCAAACGCCATTGGTATGGTAGAATTCATCTCCATTTC<br>CCGTGGGATCTACGCAACCGATCAAATGTTACAAACTTCTGATGTGGAGATCGTCACTGCTAATTCAGTCT<br>GCCCAGGAAAGTATATCGCTATCGTAAATGGGGATGTTGCGGCAGTAAAAGAAAGTGTCTCTGTAGGTGAG<br>AAGATCGCCGGAGAATTTTGGGTGGATAGCATCATTATTCCTAATGTGAACCCATTAGTGTTTCCGGCAAT<br>CACGGGAGCAACTATGCCCGACTCGATTCAGGCGTTGGGTATTATGGAGTCGTTCAGCCTGGCCACTATGG<br>TAATCGCAGCAGATGCCATTTTGAAGGCGGCAAACTTGGAGCCCCTTGATCTGCGTTTGGGGACCGGCTTG<br>GGGGGCAAGTCCTTTTTTACTTTTACAGGAGACGTAGCAGCCGTAGAGGCGGGAATTGATGCAGGGAAGTC<br>CATTGCTGAGGAAAAGGGGCTTTTGGTTAATGCAGAGGTCATTCCTTCACCATCGGATCGTCTTCTTCAGT<br>CGCTGTTGTAA |
| Cbot_hisBMC<br>TSE_K25Q | protein | MGSHHHHHHGSANAIGMVEFISISRGIYATDQMLQTSDVEIVTANSVCPGKYIAIVNGDVAAVKESVSVGE<br>KIAGEFWVDSIIIPNVNPLVFPAITGATMPDSIQALGIMESFSLATMVIAADAILKAANLEPLDLRLGTGL<br>GGKSFFTFTGDVAAVEAGIDAGKSIAEEKGLLVNAEVIPSPSDRLLQSLL* |
| Cbot_BMCTS<br>E_native_prot<br>ein_Uniprot_<br>A0A093VPU<br>0 | protein | MANAIGMVEFISISRGIYATDQMLKTSDVEIVTANSVCPGKYIAIVNGDVAAVKESVSVGEKIAGEFWVDS<br>IIIPNVNPLVFPAITGATMPDSIQALGIMESFSLATMVIAADAILKAANLEPLDLRLGTGLGGKSFFTFTG<br>DVAAVEAGIDAGKSIAEEKGLLVNAEVIPSPSDRLLQSLL* |

**Table S3. DNA and protein sequences**
